# Long-Read Sequencing of the *MUC1* VNTR: Genomic Variation, Mutational Landscape, and Its Impact on ADTKD Diagnosis and Progression

**DOI:** 10.1101/2025.09.06.673538

**Authors:** Alena Vrbacká, Anna Přistoupilová, Kendrah O Kidd, Václav Janoušek, Martin Radina, Petr Vyleťal, Ibrahim Bitar, Viktor Stránecký, Lenka Steiner-Mrázová, Helena Trešlová, Jana Sovová, Kateřina Hodaňová, Hana Hartmannová, Dita Mušálková, Klára Svojšová, Tereza Kmochová, Veronika Barešová, Abby Taylor, Lauren Martin, Antonio Sanchez, Romana Ryšavá, Innet Lajtmanová, Silvie Rajnochová-Bloudíčková, Ondřej Viklický, Gregorius Papagregorius, Constantinos Deltas, Christoforos Stavrou, Sofia Jorge, José António Lopes, Márcia Rodrigues, Elhussein Elhassan, Michelle Clince, Colm Rowan, Peter Conlon, Omri Teltsh, Gianpiero L. Cavalleri, Brendan Blumenstiel, Diana Toledo, Marina DiStefano, Matthew DeFelice, Martina Živná, Anthony J Bleyer, Stanislav Kmoch

## Abstract

**Background:** ADTKD-*MUC1* is caused by frameshift mutations in *MUC1* gene that produce a frameshifted protein (MUC1fs) toxic to kidney cells. The gene’s variable number of tandem repeats (VNTR), with high GC content, makes it largely inaccessible to standard sequencing. As a result, both the reference sequence and natural variation in this region remain poorly defined, complicating mutation detection and data interpretation. Standard methods also fail to pinpoint the exact VNTR unit affected, limiting insight into mutation mechanisms and genotype–phenotype correlations.

**Methods:** We employed Single Molecule, Real-Time (SMRT) sequencing and characterized the genomic sequence of *MUC1* in 300 individuals including 279 individuals from 143 families suspected of having ADTKD-*MUC1*. We compared these results to those obtained using the CLIA-approved mass spectrometry-based probe extension (PE) assay, which specifically detect the most prevalent 59dupC mutation. We correlated the structural features of the *MUC1* VNTR with the rate of kidney function decline in affected individuals.

**Results:** We identified *MUC1* consensus sequences for 205 unique VNTR alleles, with 9 distinct types of frameshift mutations present on 52 distinct mutated VNTR alleles. *MUC1* frameshift mutations were identified in 71 of 143 families (50%) with suspected ADTKD, comprising 135 genetically affected individuals (48%). The SMRT assay exhibited complete concordance and revealed that the PE assay is capable of detecting frameshift mutations in approximately 85% of affected families. The constellation of VNTR structures supports a genotype–progression model, in which fast progressors exhibit a significantly lower number of repeat units on the wild-type allele and a higher number of repeats on the mutation-bearing allele, including an increased number of frameshifted repeat units.

**Conclusions:** SMRT sequencing outperforms current diagnostic methods for ADTKD-*MUC1* and reveals the prognostic value of VNTR structures. Although their contribution to disease progression is modest (∼6% variance explained), it remains biologically and clinically meaningful.

**Key Points 3:**

- Single Molecule, Real-Time (SMRT) sequencing of *MUC1* outperforms existing genetic and immunohistochemical methods for ADTKD-*MUC1* diagnosis.
- CLIA-approved mass spectrometry-based genotyping assay, complemented by SMRT sequencing, can detect nearly all ADTKD-*MUC1* patients.
- Disease progression correlates with VNTR pattern: fast progressors have fewer WT repeats and more mutant/frameshifted repeats

## Introduction

Autosomal dominant tubulo-interstitial kidney disease (ADTKD) refers to a group of genetic conditions characterized by slowly progressive chronic kidney disease with a bland urinary sediment, typically leading to kidney failure between the ages 20 and 70 years [1-3]. ADTKD is caused by mutations in several genes including *MUC1 [4], UMOD [5], HNF1B [6], REN [7], SEC61A1 [8], APOA1* [9], and *APOA4* [10]; for a recent review see ref [11]. Unifying terminology using a subclassification of ADTKD according to the particular genetic defect has been proposed by Kidney Disease Improving Global Outcomes (KDIGO) [2] and globally adopted.

ADTKD-*MUC1* is caused by deletions, insertions and duplications within *MUC1* that introduce a +1 frameshift during translation. This specific frameshift leads to the production of an abnormal protein known as frameshifted MUC1 (MUC1fs), which is toxic to kidney cells [12]. Notably, only mutations that cause this specific +1 frameshift leading to the synthesis of the MUC1fs protein have been found to cause ADTKD-*MUC1*.

*MUC1* is located on chromosome 1q21. The gene contains a characteristic segment composed of variable number of tandem repeat (VNTR) units, each consisting of a degenerate 60–base pair motif repeated 20 to 125 times. Each repeat unit encodes a 20-amino acid peptide. Due to genetic polymorphisms, individual repeat units exhibit slightly different peptide sequences, leading to extensive variation in the amino acid composition the VNTR region of the MUC1 protein [13]. In addition to this extensive sequence polymorphism, there are also single nucleotide polymorphisms (e.g. rs4072037) that contribute to alternative splicing and production of multiple *MUC1* mRNA isoforms [14-16]. Thus, the *MUC1* VNTR region poses significant challenges for genetic analysis due to its highly repetitive polymorphic structure as well as its extremely high guanine/cytosine (GC) content (>80%). These features render it largely inaccessible to standard Sanger sequencing and Illumina short-read sequencing methods. As a result, the reference sequence of this region, as well as its natural genetic variation, remain poorly characterized—further complicating the interpretation of sequence data and the detection of pathogenic frameshift variants.

In other words, due to the technical limitations in analyzing the *MUC1* VNTR region, mutations causing ADTKD-*MUC1* are not detected by diagnostic multi-gene panels, whole-exome sequencing (WES), or whole-genome sequencing (WGS). Currently, these pathogenic variants can only be accurately identified at specialized research or diagnostic centers using advanced techniques specifically designed to interrogate this challenging genomic region.

Recent advances in genetic and immunohistochemical diagnostics [17-21] have enabled identification of ADTKD-*MUC1* in >1000 individuals from >320 families ([22] and unpublished). These studies revealed remarkable intra- and inter-familial variability in CKD progression [23-26] and showed that most mutations arise from single cytosine insertion into a homopolymeric tract of seven cytosines within one of the *MUC1* VNTR repeat units. This mutation, originally referred as the +C insertion or insC [4], is now designated as 59dupC, indicating that the cytosine duplication occurs in the heptanucleotide cytosine tract ending at the 59th nucleotide position of the canonical VNTR unit. Other insertions, deletions and duplications can also produce a +1 frameshift and pathogenic MUC1fs protein [27, 28]. In some cases, MUC1fs staining in urinary cells or tissue is observed despite negative genetic testing for mutations in *MUC1* and other ADTKD genes [18].

Several methods are available for *MUC1* genetic testing. In the U.S., CLIA-approved testing is performed at the Broad Institute using a mass spectrometry-based probe-extension assay [17]. This method detects the common 59dupC mutation and the 60dupA mutation [18] the later not clinically validated and reported without clinical interpretation. A fluorescence-based version of this assay is now offered by several European laboratories [29]. Additional frameshift mutations, including 59dupC, can be identified with Illumina short-read sequencing and specialized bioinformatics pipelines such as VNTyper [18, 20, 30], though the specificity and sensitivity of these approaches remain to be validated.

Thus, a key limitation in ADTKD-*MUC1* genetic diagnosis is the limited availability of testing methods and their inability to detect the full spectrum of pathogenic VNTR mutations. Moreover, none can pinpoint the exact VNTR unit harboring the frameshift mutation, which may be relevant for elucidating mutation mechanisms and genotype-phenotype correlations.

To address these limitations, we adapted the protocol of Wenzel et al, [31] employing Single Molecule, Real-Time (SMRT) sequencing, with a custom analysis pipeline, to characterize the *MUC1* genomic sequence in 300 individuals including 279 individuals from 143 families with suspected ADTKD-*MUC1*. We assessed VNTR length, repeat structure, and frameshift mutations, and compared findings with the CLIA-approved mass spectrometry–based assay. We further correlated mutation location with kidney failure onset and progression. Our results demonstrate the superiority of SMRT sequencing over existing diagnostic methods and highlight its potential for prognostic stratification and personalized management in ADTKD-MUC1.

## Methods

### Ethical approval

This study was approved by the Institutional Review Boards of the First Faculty of Medicine, Charles University in Prague and the Wake Forest University Health Science Institutional Review Board.

### Clinical evaluation and study population

The Wake Forest ADTKD registry is comprised of over 1000 families referred to A.J.B. by physicians and/or family members since 1996. Families are screened by A.J.B. and the following data is collected: demographics, pedigree, medical history, laboratory values, imaging results, and biopsy reports.

### Mass Spectrometry-Based Assay for the Molecular Diagnosis of ADTKD-*MUC1*

A probe (primer) extension assay that in principle detects only mutations that are destroying MwoI restriction site within mutated VNTR was performed using a CLIA-approved mass spectrometry-based assay at the Broad Institute of MIT and Harvard (Cambridge, MA) [17, 18].

### Long range PCR amplification and Long-read sequencing of *MUC1*

Whole blood DNA was extracted using standard methods, and the *MUC1* VNTR was PCR-amplified as described[18]. Libraries were sequenced on the PacBio Sequel I system, CCS reads were generated with SMRT Link v8.0, and analyzed with the PacMUC1 pipeline (**Figure 1**).

**Figure 1.**
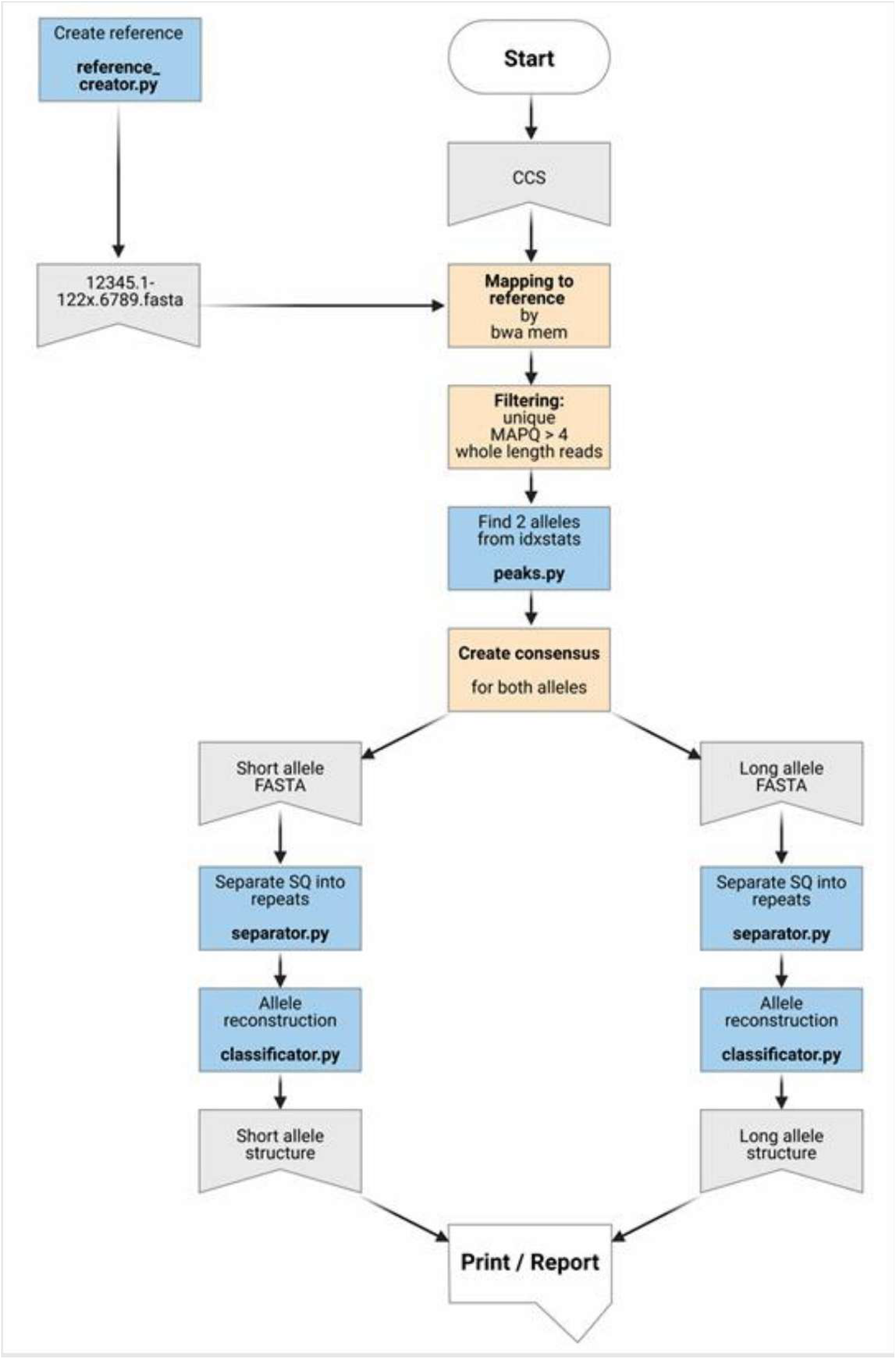
The PacMUC1 Bioinformatics pipeline. First, a reference sequence of MUC1 was created that contained 120 contigs, with each contig containing pre-repeats (1, 2, 3, 4, 5) and after-repeats (6, 7, 8, 9), separated by 1-120 canonical × repeats. The CCS FASTQ files were mapped to the reference sequence using bwa-mem (version 0.7.16a-r1181) and postprocessed using samtools (version 1.9) [40]. Chimeric, incomplete, and low quality reads were removed, and the aligned sequences were displayed in the Integrative Genomics Viewer (IGV, version 2.16.2) for visual inspection. To determine the length of each of the two alleles for each sample, idxstats were generated using samtools (version 1.9) and peaks with the highest number of mapped reads were selected. To find changes f rom the reference sequence we used the deep neural network-based variant caller Clair (version 2.0. 7), https://github.com/HKU-BAL/Clair and Clair3 (version 1.0.10), https://github.com/HKU-BAL/Clair3 with a model trained for Pac Bio data. The resulting VCF file and bcftools (version 1.10.2) were used to create a consensus sequence for each of the determined alleles. Custom Python scripts were used to separate the consensus sequences into individual repeats, classify them (each unique 60-base repeat segment is represented by a different letter or number, corresponding to the repeats previously identified by Kirby et al. [4] and Wenzel et al. [31], calculate statistics, and generate reports. The obtained MUC1 VNTR sequences are reported as a sequence of unique repetitive units, characterized by numbers (pre- and after-repeats) and letters (canonical repeats), as originally described by Kirby et al. **[4]. A** minimum read coverage of 1 Ox was required for reliable determination of the VNTR allele structure. In addition, for related individuals, only a single representative allele was reported to prevent artificially inflated allele occurrence or frequency. A minimum read coverage of 1 Ox was required also for reliable identification of the frameshift mutation. Cases in which the mutation was clearly identified (eg. visible in the IGV) but supported by fewer than 1 Ox reads were reported as inconclusive. Created with BioRender.com

### Nomenclature of the *MUC1* VNTR sequence and frameshift mutations

The *MUC1* VNTR comprises 60-base pair repeats encoding 20–amino acid peptides; **Figure 2A** shows standardized nomenclature for length variants, repeat structures, and frameshifts.

**Figure 2.**
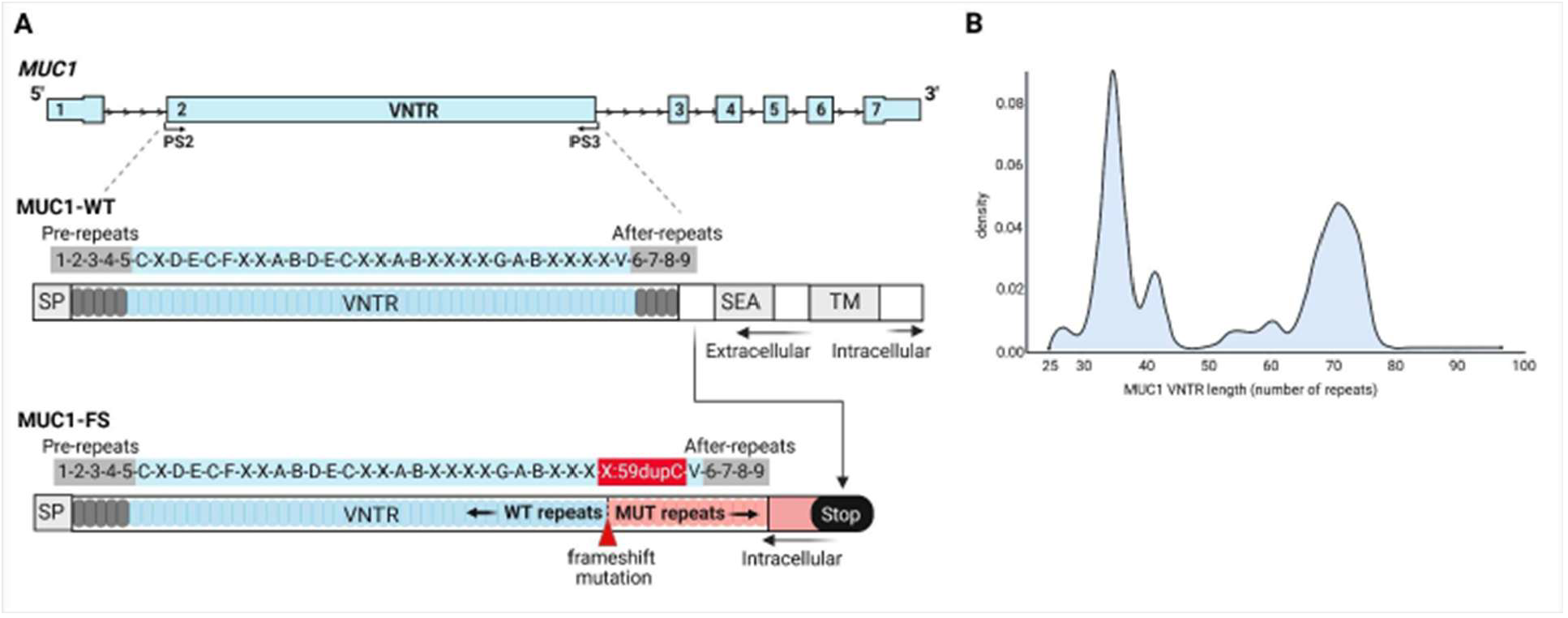
*MUCl* VNTR structure. **(A)** The canonical sequence of *MUC1* consists of exon 1, which encodes a conserved N-terminal signal peptide (SP); exon 2, which encodes the VNTR region; and exons 3-7, which encode the conserved C-terminal domains, including the SEA (Sperm protein, Enterokinase, and Agrin) domain and transmembrane domain (TD), both of which are critical for protein stability, trafficking, and function. The sequence of the *MUC1* VNTR is reported as originally described by Kirby et. al ^4^, with the following characteristics: There are 5 pre-repeats prior to theVNTR, each designated with the number 1-5 respectively. Variants in these pre-repeats are followed by an apostrophe. For example, 3’ references a third pre-repeat with a variant within it. The letters A-Z and aA, aB - aZ, bA-bZ, cA-cZ … and beyond, defining the sequence of individual types of canonical repeats and the numbers 6-9 defining individual types of after-repeats (including their variants labeled with an apostrophe, “‘“, for example 6’). For example, the sequence 1-2-3-4-5-C-X-D-E-C-F-X-X-A-8-D-E-C-X-X-A-8-X-x-x-x-G-A-8-X-X-X-X-X-X-V-6-7-8-9 defines the VNTR that is composed of the 5 pre-repeats, 29 canonical repeats of different types and 4 after-repeats. In the case of long-read sequencing, frameshift mutations can be assigned to a precise localization within the VNTR allowing exact mapping of the mutation within the full repeat sequence and enabling detailed sequence comparison. For example, in this case, the sequence indicates the presence of the 59dupC mutation in the 28th VNTR repeat unit, designated as unit X. This mutation leads to the production of an abnormal protein known as frameshifted MUC1 (MUC1fs), which consists of an initial wild-type *MUC1* sequence (blue) followed by a mutation-induced frameshifted segment containing mutated VNTR units. The translational frameshift introduces a premature stop codon shortly after the VNTR region, resulting in the loss of the characteristic SEA domain and transmembrane domain. As a consequence, MUC1fs fails to localize to the cell membrane, remains intracellular, and exhibits toxic effects on kidney cells. Novel synonymous and missense variants and in-frame insertion or deletion variants defining new types of pre-repeats, canonical repeats and after-repeats are described as above in ascending order either with new numbers or letters. **(B)** Distribution and frequencies of VNTR lengths, defined by the number of repeat units, across 598 alleles. Created with BioRender.com

### Statistical analysis

Associations between eGFR decline and allele repeat lengths were tested using Spearman’s correlation and linear regression. Sequential models assessed the incremental contribution of WT and mutated allele lengths, with significance set at p < 0.05.

Methodological details are available in the **Supplementary Appendix**.

## Results

### Natural genomic variation of *MUC1* VNTR

We analyzed *MUC1* VNTR sequences in 300 individuals, including 21 healthy volunteers and 279 subjects from 143 ADTKD families. VNTR lengths were determined for 557 of 600 expected alleles (93%), with allele length distribution and frequencies shown in **Figure 2B**.

We established the consensus sequence for both VNTR alleles in 202 (67%) individuals. In 98 individuals (30%), we were able to establish the consensus sequence for only the shorter allele, likely due to significant differences in allele lengths and long range PCR-inherent allelic dropout, which resulted in minimum read coverage below the 10× threshold required for reliable determination of the VNTR allele structure. We identified consensus sequences for 502 VNTR alleles, comprising 205 unique alleles (41%) (**Supplementary Table 1**). In total, 78 distinct repeat units were detected, 44 of them novel (56%): 9 pre-repeat units (2 novel), 63 canonical units (41 novel, including one with an in-frame 18 bp indel), and 6 after-repeat units (1 novel). The most common units were × (49%), A (13%), B (9%), and C (4%). An updated list of repeats is available at https://github.com/pristanna/muc1repeats. Researchers may submit novel VNTR allele and repeat unit sequences via this link. All identified VNTR repeat units, labeled by one-letter codes following Kirby et al.[4] and Wenzel et al.[31] and extended in this study, are shown in **Figure 3**. Pre- and after-repeat unit sequences are provided in **Supplementary Table 2**.

**Figure 3.**
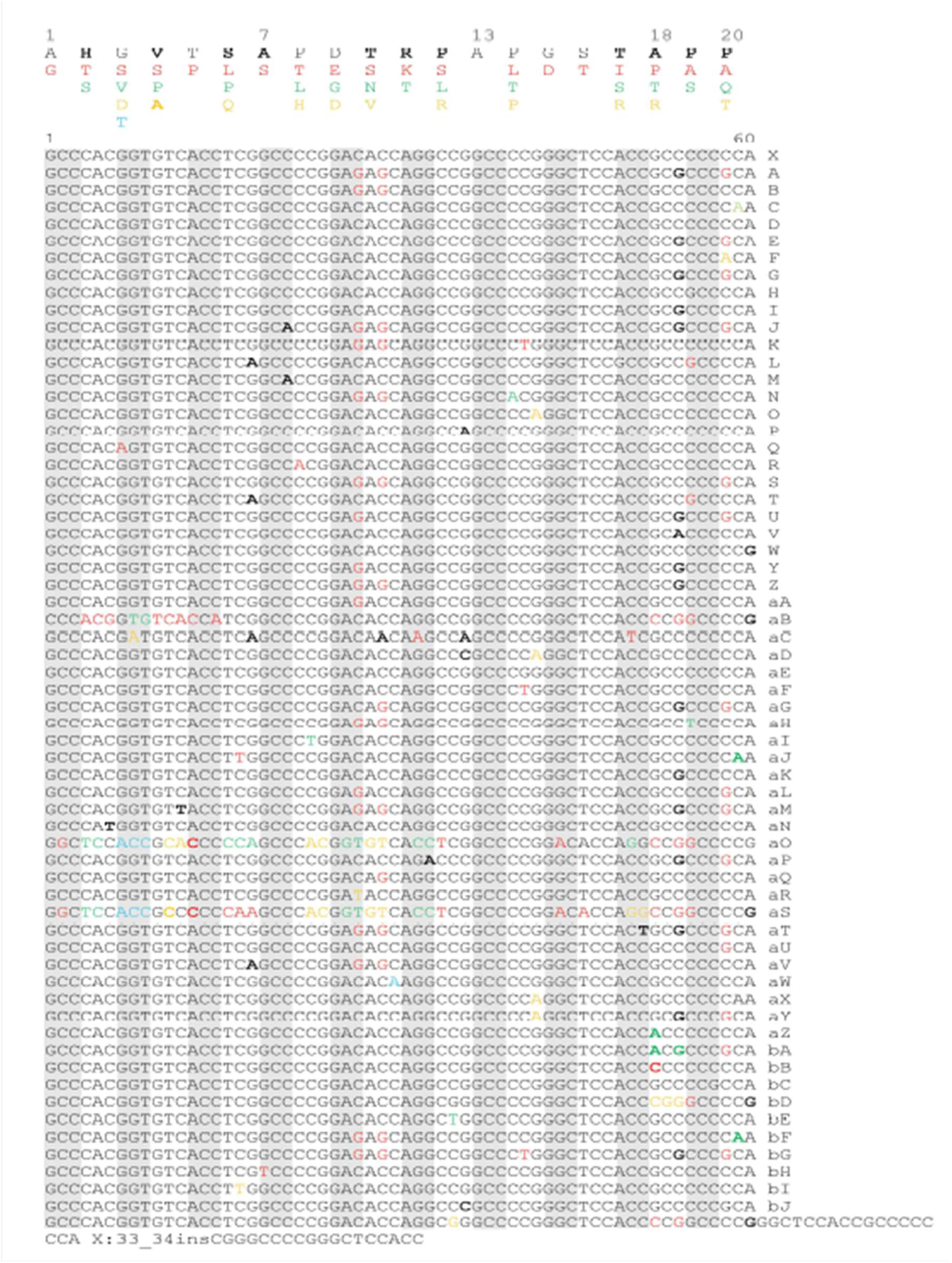
VNTR repeat sequences. The list of all V TR repeat sequences identified and labeled with one letter codes according to Kirby et al.^4^ and Wenzel et al.^28^, and extended in this study. A 60-nucleotide sequence of the canonical VNTR repeat u it ‘X’ is shown first, followed by nucleotide sequences of alternative VNTR units labeled with letters. synonymous sequence variants are shown in bold black font. Missense variants are highlighted in colors corresponding to amino acid substitutions within the canonical 20-amino acid VNTR unit that is shown at the top. Note a high degree of conservation of four alanine residues {Al, A7, A13 and A18), which are regularly spaced throughout each repeat unit.

### *MUC1* frameshift mutations identified in the Wake Forest ADTKD registry

SMRT sequencing was performed in 279 individuals from 143 families suspected of ADTKD-*MUC1*, including 144 clinically affected individuals with an eGFR of less than 60 mL/min/1.73 m^2^, and 135 clinically unaffected individuals with normal eGFR. In this cohort, the MALDI-TOF probe extension (PE) genotyping for 59dupC or 60dupA mutations (detected as analytes at 5904 Da or 6571 Da, respectively) was performed at the Broad Institute [17, 18]. **Table 1** summarizes PE genotyping in 202 individuals from 128 families. Pathogenic variants were detected in 59 families (109 individuals): 58 with 59dupC and 1 with 60dupA. Two families had inconclusive results—one with a weak 59dupC signal and one with a novel 6883 Da analyte of uncertain significance (**Figure 4**). The remaining 67 families (91 individuals) tested negative (wild-type).

**Table 1.**
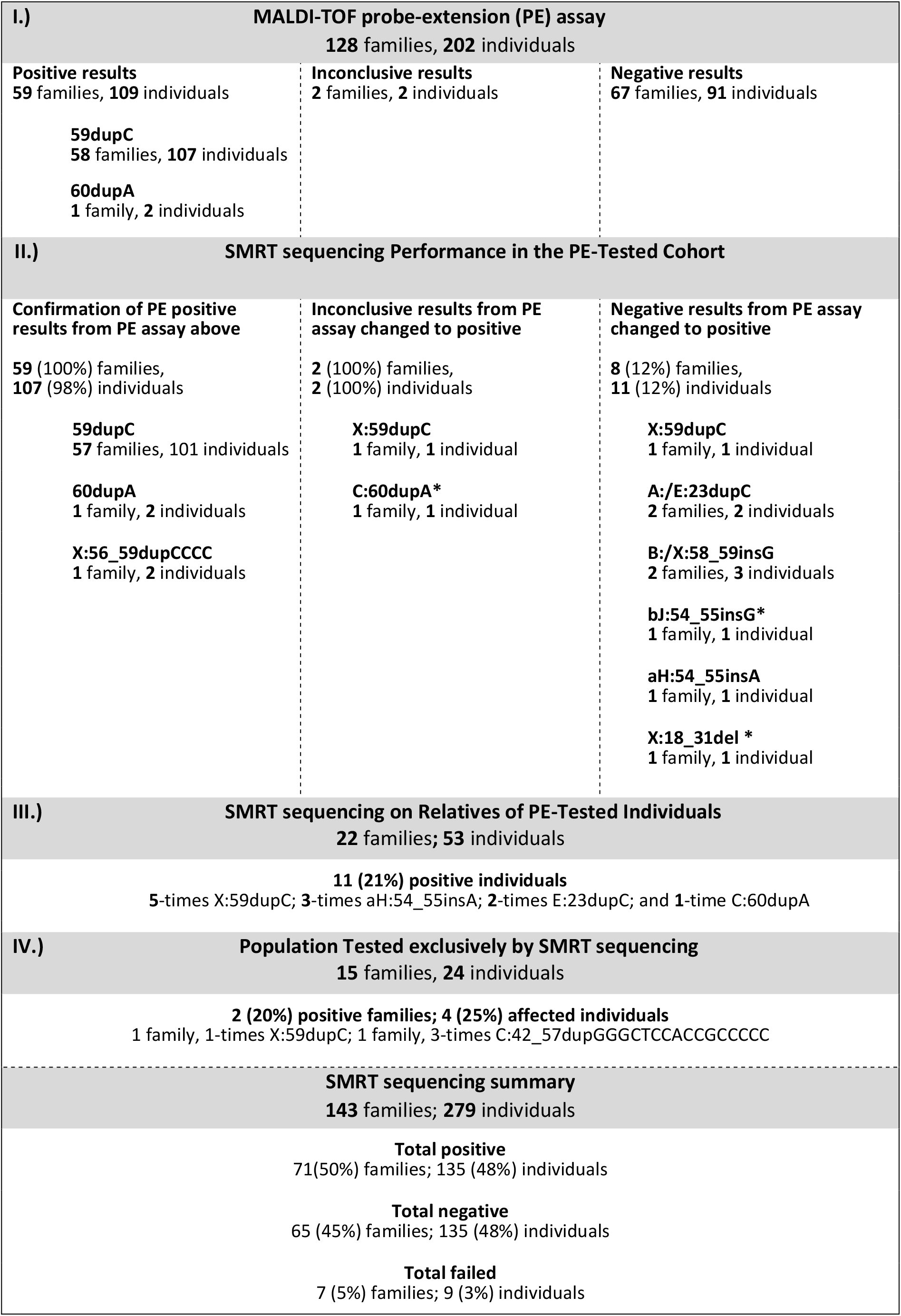
MUC1 frameshift mutations identified in the Wake Forest ADTKD registry.

**Figure 4.**
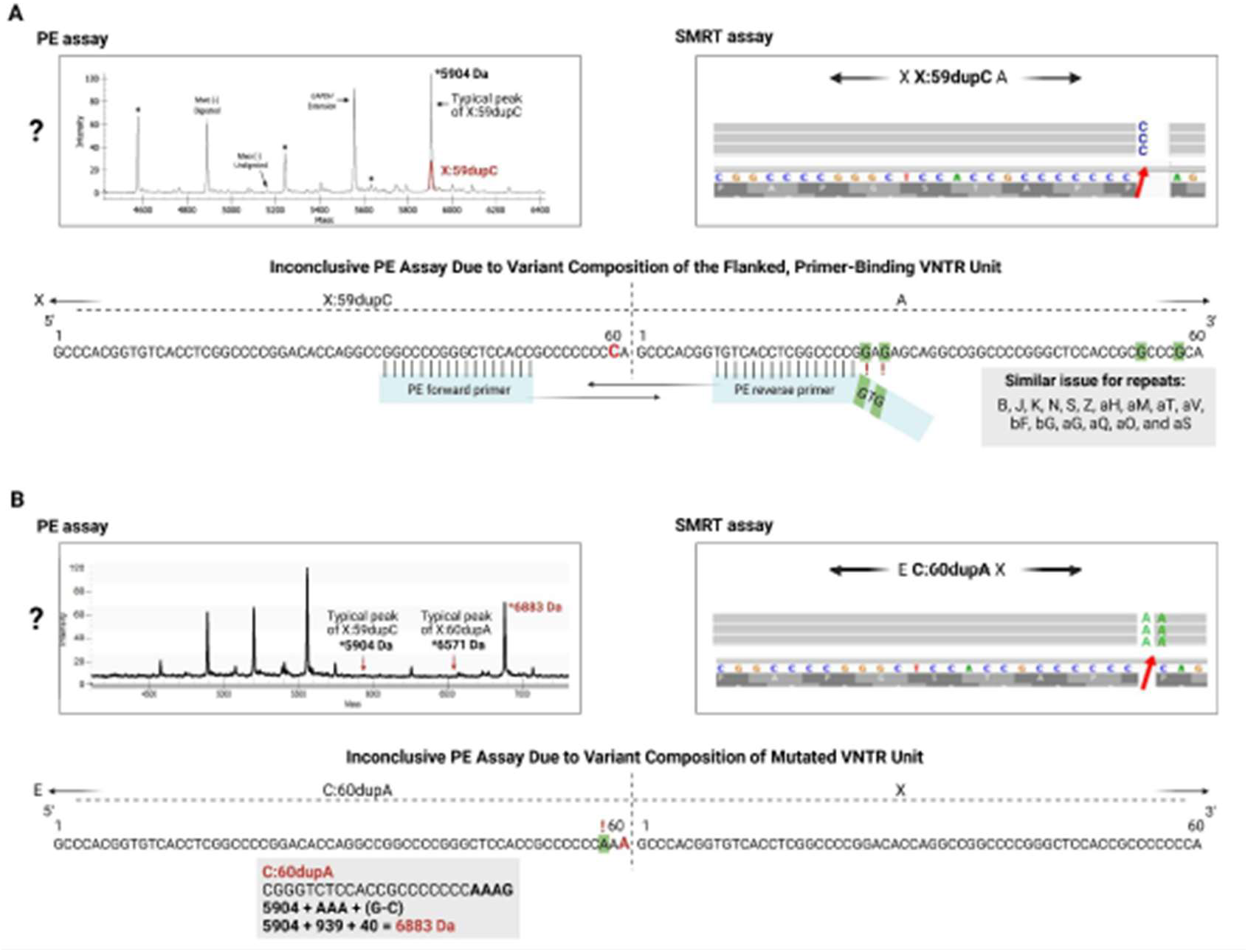
VNTR structures in cases with inconclusive results from the PE genotyping assay. {**A**) Inconclusive PE assay due to variant composition of the flanked, primer-binding VNTR unit. In this case, the PE assay identified an extension analyte at 5904 Da, indicating the presence of the 59dupC, but with a low signal-to-noise ratio {red peak). The SMRT assay identified 59dupC mutation located within the repeat unit X, which is flanked upstream by the repeat unit A, forming the haplotype XA. The sequence of the repeat unit A is not fully complementary to the tailed primer nested within its 60-bp sequence, which negatively affects the initial PCR amplification of the Mwol-intact VNTR fragments. Notably, similar PCR amplification issues during the PE assay may occur in hypothetical cases where the mutated repeat unit is followed by specific repeat units such as B, J, K, N, S, Z, aH, aM, aT, av, bf, and bG. Additional repeat units, including aG, aQ, aO, and aS, may also influence the PE assay in a similar manner. (**B**) Inconclusive PE assay due to variant composition of mutated VNTR unit. In the second case, the PE assay identified a specific extension analyte at 6883 Da, whose origin and diagnostic significance remained unknown. The SMRT assay revealed that this signal corresponds to an adenine duplication at position 60 of repeat unit C {C:60dupA), resulting in the incorporation of an additional adenine into the primer extension product. Created with BioRender.com

In this cohort, the SMRT assay showed 100% concordance with the PE assay at the family level and 98% at the individual level, detecting mutations in all 59 families and 107/109 individuals positive by PE. One individual initially reported as 59dupC was reclassified as X:56_59dupCCCC. In two PE-positive cases, SMRT assay failed due to allelic dropout of large (68–74 units) alleles paired with shorter (34–41 units) alleles. For the two PE-inconclusive cases, SMRT identified X:59dupC in one and C:60dupA in the other (**Figure 4**).

Among 67 families (91 individuals) with a negative (wild-type) results of PE genotyping assay, the SMRT assay identified an additional 8 families (12%) with 11 individuals (12 %) harboring 8 distinct types of frameshift mutations. In one instance, for the same reason observed in an inconclusive case with the XXA haplotype (see **Figure 4**), the SMRT assay identified the X:59dupC mutation, leading to a 1% discordance rate among all cases that tested negative with the PE assay. The other seven types of frameshift mutations could not be detected by the PE assay. This limitation was due to the specific positions of the individual mutations within their respective repeat units.

These mutations either do not disrupt the MwoI restriction site, which cleaves within the 5′-GCNNNNN/NNGC-3′ sequence of the 60-bp repeat unit - a critical step for the specific PCR amplification of the mutated repeat unit in the PE assay, or they alter the VNTR sequence complementary to the assay primer, thereby limiting the primer extension reaction and assay’s ability to detect them.

The SMRT assay was also performed in 77 individuals who had not been previously genotyped using the PE assay. Among 53 individuals from 22 families with known mutations (identified earlier by either the PE or SMRT assay), it identified 11 additional genetically affected individuals. In addition, the SMRT assay identified 4 genetically affected individuals in 2 (17%) of the 15 previously unsolved families carrying either X:59dupC or C:42_57dupGGGCTCCACCGCCCCC mutation.

Thus, the SMRT assay identified *MUC1* frameshift mutations in 71 of 143 families (50%) with suspected ADTKD, involving 135 genetically affected individuals (48%). In an additional 133 individuals (also 48%), both *MUC1* alleles were successfully sequenced but no mutation was detected. The assay failed in 9 individuals (3%) from 7 families, in whom only one *MUC1* allele was successfully sequenced—likely due to allelic dropout, a known limitation of long-range PCR and SMRT sequencing.

### Concordance of the MUC1fs genotyping/sequencing with immunohistochemical testing for MUC1fs

A noninvasive immunohistochemical method to detect MUC1fs on various epithelial tissues and urinary cell smears proved to be useful for diagnostic testing of ADTKD-*MUC1* [18, 19]. Assuming high specificity and sensitivity, we used the SMRT assay results as the definitive reference, allowing a more accurate assessment of the sensitivity and specificity of the immunohistochemical stainings. Here, renal biopsies were available from 21 clinically affected individuals with 18 positive, 1 inconclusive and 2 negative immunohistochemical stainings for MUC1fs. Among the 18 individuals with positively stained kidney specimens, the SMRT assay identified one of the known frameshift mutations in 16 cases (89%), while no mutation was detected in the remaining 2 individuals (11%). Of these 16 individuals who tested positive by the SMRT assay, 4 (25%) harbored frameshift mutations that could not be detected by the PE assay. In one individual with an inconclusively stained specimen, neither the PE assay nor the SMRT assay identified frameshift mutation. Among two individuals with negative immunohistochemical staining, one was found with the X:59dupC mutation, while the other tested negative by both assay types. Thus, positive immunohistochemical staining of kidney tissue for MUC1fs is highly diagnostic for ADTKD-*MUC1*; however, negative staining does not exclude the diagnosis.

In addition, we had urinary smears from 78 individuals with 46 positive, 6 inconclusive and 26 negative immunohistochemical stainings for MUC1fs. Among the 46 individuals with positively stained specimens, the SMRT assay identified one of the known frameshift mutations in 35 cases (76%), while no mutation was identified in the remaining 11 individuals (24%). Of these 35 individuals who tested positively by the SMRT assay, 30 (86%) harboured the X:59dupC mutations, 5 (14%) harbored frameshift mutations that could not be detected by the PE assay. Among 6 individuals with inconclusive immunohistochemical MUC1fs stainings there were 2 individuals (33%) with the X:59dupC mutation identified by both assay types and 4 individuals (66%) who were negative for frameshift mutation by both assay types. Among 26 individuals with negative immunohistochemical MUC1fs staining, 3 (12%) were found to carry frameshift mutations: 2 individuals with the X:59dupC mutation identified by both assays, and 1 individual with the X:58_59insG mutation detected only by the SMRT assay. The remaining 23 individuals tested negative for frameshift mutations by both assay types. Thus, positive immunohistochemical staining of urinary cells for MUC1fs is also highly diagnostic for ADTKD-*MUC1*; however, it shows lower sensitivity and specificity compared to kidney tissue. As with kidney tissue, negative staining does not exclude the diagnosis.

Thus, in the course of our work, we identified 9 types of frameshift mutations (**Table 1 and Supplementary Table 3**) present on 52 distinct mutated VNTR alleles (**Supplementary Table 4**), with some mutated VNTR alleles shared among reportedly unrelated families. The most frequent mutation was 59dupC identified on 42 VNTR alleles (81%), while the other eight mutations occurred at most twice. The mutational spectrum indicated that the PE assay may detect frameshift mutations in 44 VNTR alleles, representing approximately 85% of affected families.

### The relation of the *MUC1* VNTR structure to the age of onset and rate of kidney disease progression

The length of the *MUC1* VNTR has been positively associated with several kidney-related traits, including blood urea nitrogen and serum urate levels, and an increased risk of gout and chronic tubulointersticial nephritis [32, 33]. Individuals with the *MUC1* mutation suffer from chronic kidney disease with a widely variable age of onset of kidney failure ranging from 16 to >80 years [23]. To this end, we have used sequence information and correlated individual wild-type and mutated VNTR structures with the age of onset of kidney failure and rate of kidney disease progression in individuals with MUC1fs mutations.

The average age of kidney failure in all ADTKD-*MUC1* patients in our registry (n=447) is 43.46 ±14.0 years; and for the WF VNTR Cohort with VNTR length determination (n=268) 44.9 ± 14.3 years (t-test P = 0.19, indicating no bias in sampling). To calculate the rate of eGFR decline over time, we assumed an eGFR at birth of 120 ml/min and calculated eGFR decline as (eGFR-120)/age (years). For those who reached kidney failure, their eGFR was defined as 10 mL/min at kidney failure. If decline was ≤ -2.5ml/min/year, the patient was defined as a fast progressor, and if > -2.5 ml/min/year as a slow progressor. Progression status (fast vs. slow) along with VNTR structure information was available for 149 genetically affected individuals, comprising 65 males and 84 females (**Supplementary Table 5)**.

Fast progressors, compared to slow progressors, had a significantly lower number of repeat units on the wild-type allele (median: 37 vs. 57; one-tailed Mann–Whitney *p* = 0.01828) and a higher number of repeat units on the frameshift mutation-harboring allele (median: 69 vs. 59; one-tailed Mann–Whitney *p* = 0.01958). Consequently, fast progressors, compared to slow progressors, exhibited a higher number of both wild-type repeat units (median: 15 vs. 9; one-tailed Mann–Whitney *p* = 0.03583) and frameshifted repeat units (median: 46 vs. 35; one-tailed Mann–Whitney *p* = 0.00318) on the mutated allele (**Figure 5**). This supports an additive genotype–progression model, with eGFR decline correlating with the repeat length difference between wild-type and mutated alleles (Spearman’s ρ = 0.235, p = 0.004; ρ^2^ ≈ 5.5%). Linear regression confirmed that larger mutated allele length predicted faster decline (β = –0.024, p = 0.029), while WT allele length showed only a trend (β = 0.019, p = 0.054). The model explained ∼6% of the variance, indicating a modest but biologically meaningful contribution of allele length variation.

**Figure 5.**
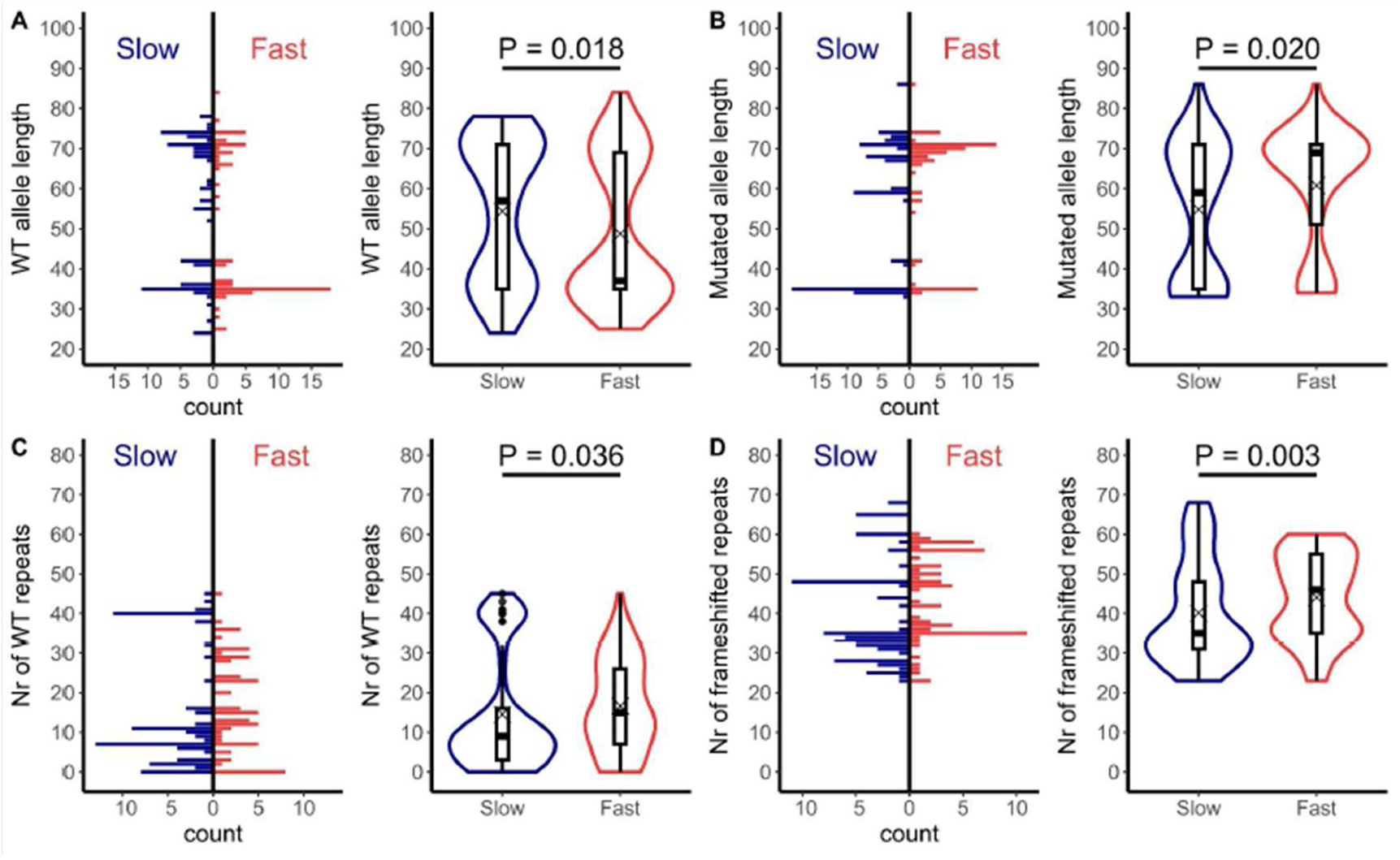
MUCl VNTR structure and the age of onset and rate of kidney disease progression. Distribution and frequencies of VNTR lengths were defined by the number of repeat units. The rate of eGFR decline over time was calculated as eGFR-120/age (years). For those who reached ESRD, their eGFR was defined as 10 ml/min at their age ESRD. If decline was ≤ -2.Sml/min/year, the patient was defined as a fast progressor (1; red), and if > -2.5 ml/min/year as a slow progressor (0; blue). Fast progressors, compared to slow progressors, had (**A**) a significantly lower number of repeat units on the wild-type (WT) allele (median: 37 vs. 57; one-tailed Mann-Whitney p = 0.01828) and (**B**) a higher number of repeat units on the frameshift mutation-harboring allele (median: 69 vs. 59; one-tailed Mann-Whitney p = 0.01958). Consequently, fast progressors, compared to slow progressors, exhibited (**C, D**) a higher number of both wild-type repeat units (median: 15 vs. 9; one-tai led Mann-Whitney p = 0.03583) and frameshifted repeat units (median: 46 vs. 35; one-tailed Mann-Whitney p = 0.00318) on the mutated allele.

## Discussion

The repetitive, GC-rich nature of the MUC1 genomic sequence makes it difficult to analyze, leaving reference sequences and natural variation poorly characterized. In this study, we applied SMRT sequencing with a custom pipeline to assess *MUC1* variation, including VNTR length and unit structure, in 279 individuals from 137 families suspected of ADTKD-*MUC1* (144 affected, 135 unaffected). Our goals were fourfold: (i) characterize natural VNTR variation, (ii) identify pathogenic frameshift mutations for genetic diagnosis, (iii) evaluate the diagnostic performance of the CLIA-approved mass spectrometry assay performed at the Broad Institute of MIT and Harvard [17] and MUC1fs immunohistochemistry in patient-derived materials performed at the First Faculty of Medicine, Charles University, Prague [18], and (iv) investigate the prognostic relevance of VNTR structures for variability in disease onset and progression.

### Methodology

The SMRT assay methodology we employed has limitations inherent to both long-range PCR and SMRT sequencing, particularly regarding allelic dropout. Suboptimal DNA quality, combined with substantial differences in *MUC1* allele lengths may result in incomplete representation—here defined as <10x read coverage—or complete absence of the longer allele. If needed for diagnostic purposes, these technical limitations could be resolved in individual cases through alternative DNA isolation procedures to prevent DNA damage, optimization of the PCR protocol, and increased sequencing coverage. An alternative approach not explored in this study may represent an amplification-free protocol for targeted enrichment of genomic regions suitable for single-molecule sequencing [34].

### Natural genetic variability

The *MUC1* gene exhibits substantial genetic variation, particularly within its VNTR region [35]. In this work we extended findings of earlier studies of Kirby et al.[4], and Wenzel et al.[31] to provide the most comprehensive list of *MUC1* VNTR sequences reported to date. Specifically, we identified consensus sequences for 205 VNTR alleles that were unique in length and composition. We identified 78 different VNTR repeat units (of which 44 were novel; (56%)). These are represented by 9 pre-repeat units (including 2 novel), 63 canonical repeat units (including 41 novel, with one of which contains in-frame 18 bp insertion/deletion), and 6 after-repeat units (including, 1 novel), see **(Figure 2 and Supplementary Table 2)**. Comparison of the VNTR sequences revealed a high degree of conservation of four alanine residues (A1, A7, A13 and A18), which are regularly spaced throughout each repeat unit. This pattern of alanine-proline motifs was observed across multiple allelic variants and suggests a functionally important role for alanine residues in maintaining the structural integrity of the VNTR domain. Given alanine’s small, non-polar side chain and its known tendency to support extended conformations without disrupting local secondary structure, the conserved spacing may contribute to the rigidity and alignment of adjacent glycosylation sites within the mucin repeat. This structural arrangement likely facilitates efficient O-glycosylation of neighboring serine and threonine residues [36], which is critical for MUC1’s function as a protective epithelial barrier [37]. The consistent presence and position of alanine across diverse repeat variants [38] further imply evolutionary selection to preserve this feature in MUC1 protein.

### Genetic diagnosis of ADTKD-MUC1

In course of our work we identified 9 types of mutations present on 52 distinct mutated VNTR alleles. The SMRT assay outperformed the PE assay by successfully identifying frameshift mutations in two families with cases that had inconclusive PE results, as well as in eight additional families comprising 11 genetically affected individuals whose mutations were undetectable by the PE assay. When sucesfull, the SMRT assay showed 100% concordance with the PE assay, except in two cases where it failed due to the technical limitations described above. The most frequent mutation was 59dupC, which was identified on 42 VNTR alleles (85%), while the other eight mutation types occurred at most twice. This finding confirms that the 59dupC mutation is the most frequent genetic alteration causing ADTKD-*MUC1*. The mutation likely arises through replication slippage, a well-described mechanism of mutagenesis that is particularly common in tandemly repeated sequences. Such sequences are prone to insertion or deletion events, and their mutation rates can be 10-to 100,000-fold higher than the average mutation rate in non-repetitive genomic regions [38, 39]. This finding has two important implications for the diagnosis of ADTKD-*MUC1*. First, it suggests that *de novo* mutations in *MUC1*-particularly the recurrent 59dupC mutation-may be more frequent than previously recognized. Both germline and post-zygotic somatic mutations should be considered in individuals with sporadic CKD, even in the absence of a family history. Second, the observed mutational spectrum indicates that the CLIA-approved mass spectrometry-based *MUC1* genotyping assay can detect frameshift mutations in approximately 85% of affected families. Therefore, for individuals who test negative using this assay but remain clinically suspicious, additional testing using SMRT sequencing and/or immunohistochemical staining for MUC1fs should be considered to improve diagnostic sensitivity.

### Prognostic potential of individual MUC1 VNTR structures

Our findings suggest that the structure of the *MUC1* VNTR—including the length and composition of both wild-type and mutant alleles—may carry prognostic significance in individuals with ADTKD-*MUC1*. Notably, fast progressors exhibited a significantly lower number of repeat units on the wild-type allele and a higher number on the mutation-bearing allele, including an increased number of frameshifted repeats. This pattern supports a genotype–progression model, as illustrated in **Figure 6**. These observations raise the possibility that individual- or family-specific VNTR configurations could serve as prognostic biomarkers, enabling risk stratification and more personalized clinical management. However, the modest effect size suggests that additional genetic and environmental modifiers contribute to the variability in disease progression. Further validation in larger, longitudinal cohorts will be necessary to confirm the predictive value of specific *MUC1* VNTR features for kidney disease progression.

**Figure 6.**
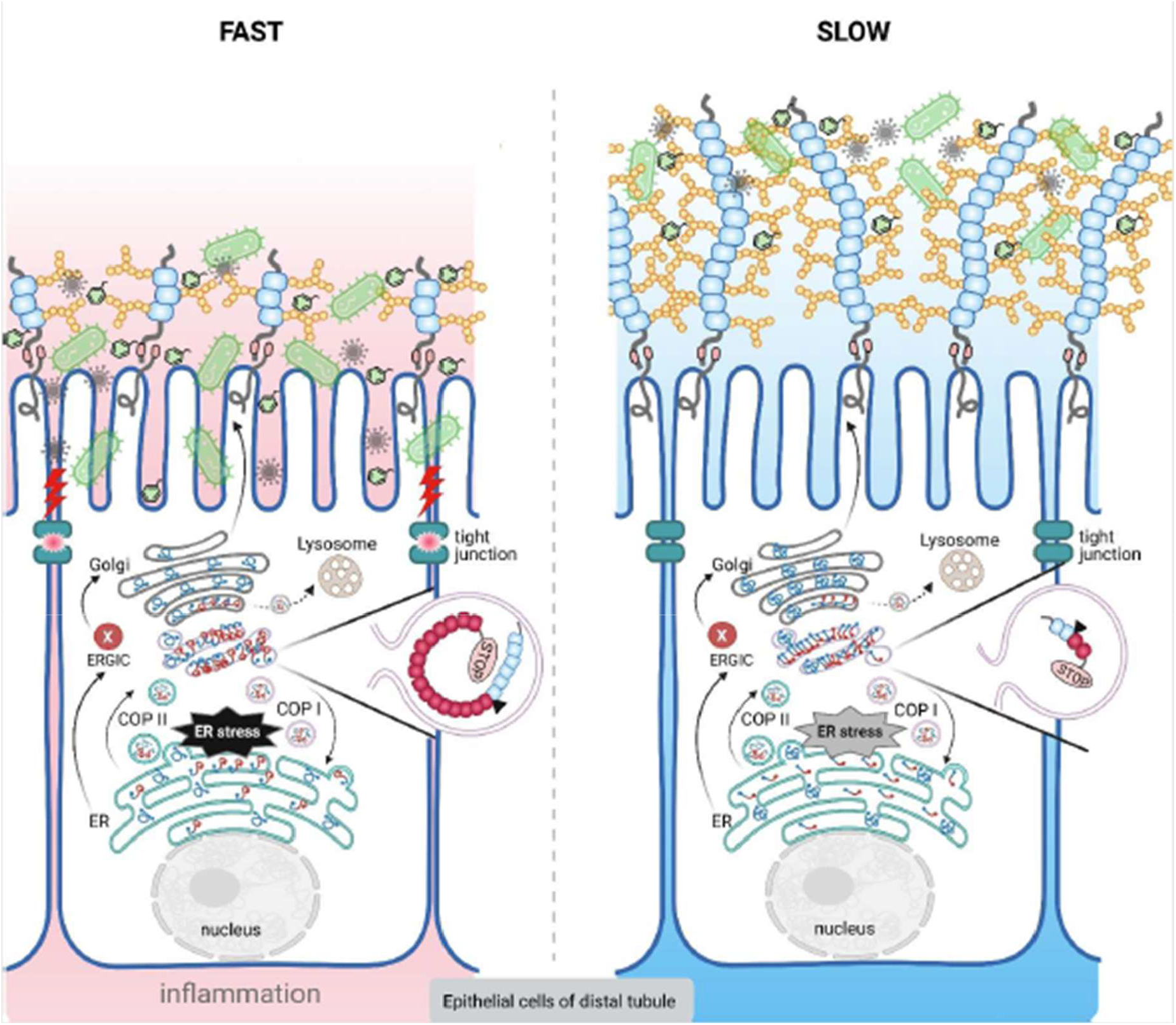
Genotype-progression model. In tubular epithelial cells of the kidney, as well as in other epithelial cells, MUC1 is synthesized in the endoplasmic reticulum (ER) and transported via COPlI-coated vesicles through the ER-Golgi intermediate compartment (ERGIC) to the Golgi apparatus, where extensive O-glycosylation (yellow circles) occurs in the tandem repeat region (blue bullets). Autoproteolytic cleavage within the SEA domain (pink bullet) generates two subunits: the extracellular MUC1-N, which is often shed and contributes to protective and signaling functions, and the membrane-tethered MUCl-C, which remains at the cell surface and participates in intracellular signaling, particularly under stress or injury. The MUClfs protein is either retrotranslocated from the ERGIC and/or Golgi to the ER via COPI vesicles for potential refolding, or directed to lysosomes for degradation. In FAST progressors compared to SLOW progressors, MUC1 haploinsufficiency—caused by intracellular retention of the mutant MUClfs protein—is further aggravated by a reduced number of tandem repeats on the wild-type allele. This may weaken the epithelial barrier along the nephron, particularly in the distal tubule and collecting duct, where MUC1 is highly expressed. Barrier disruption can lead to epithelial stress, chemical toxicity, infection, and inflammation, promoting tubular damage and fibrosis. Simultaneously, a greater number of both wild-type and frameshifted repeats (red bullets) may enhance the toxic intracellular accumulation of MUClfs protein, resulting in chronic ER stress and tubular injury. Created with BioRender.com

### Clinical testing approach

**Figure 7** shows that healthcare providers should consider using the probe-extension (PE) assay, a CLIA-approved test that is widely accessible (see www.rikd.org/resources) and identifies approximately 85% of known *MUC1* mutations. This assay represents an excellent first-line diagnostic resource. For families who test negative by the PE assay, Single Molecule, Real-Time (SMRT) sequencing can be pursued as a follow-up for more comprehensive analysis. Alternatively, depending on availability and clinical judgment, SMRT sequencing may be used as the initial diagnostic test. If the genetic testing is negative, referral to a specialized center is recommended, where immunohistochemical staining of urine samples and available kidney biopsies can be performed. Thereafter, or in parallel, whole-exome sequencing and whole-genome sequencing may be conducted to identify other potential genetic causes of ADTKD. In particular, VNTyper offers a novel bioinformatics approach for detecting *MUC1* mutations from whole-exome—and potentially in the future, whole-genome—sequencing data. It is currently under development and validation, and is expected to become a valuable tool for the genetic diagnosis of ADTKD-*MUC1*.

**Figure 7.**
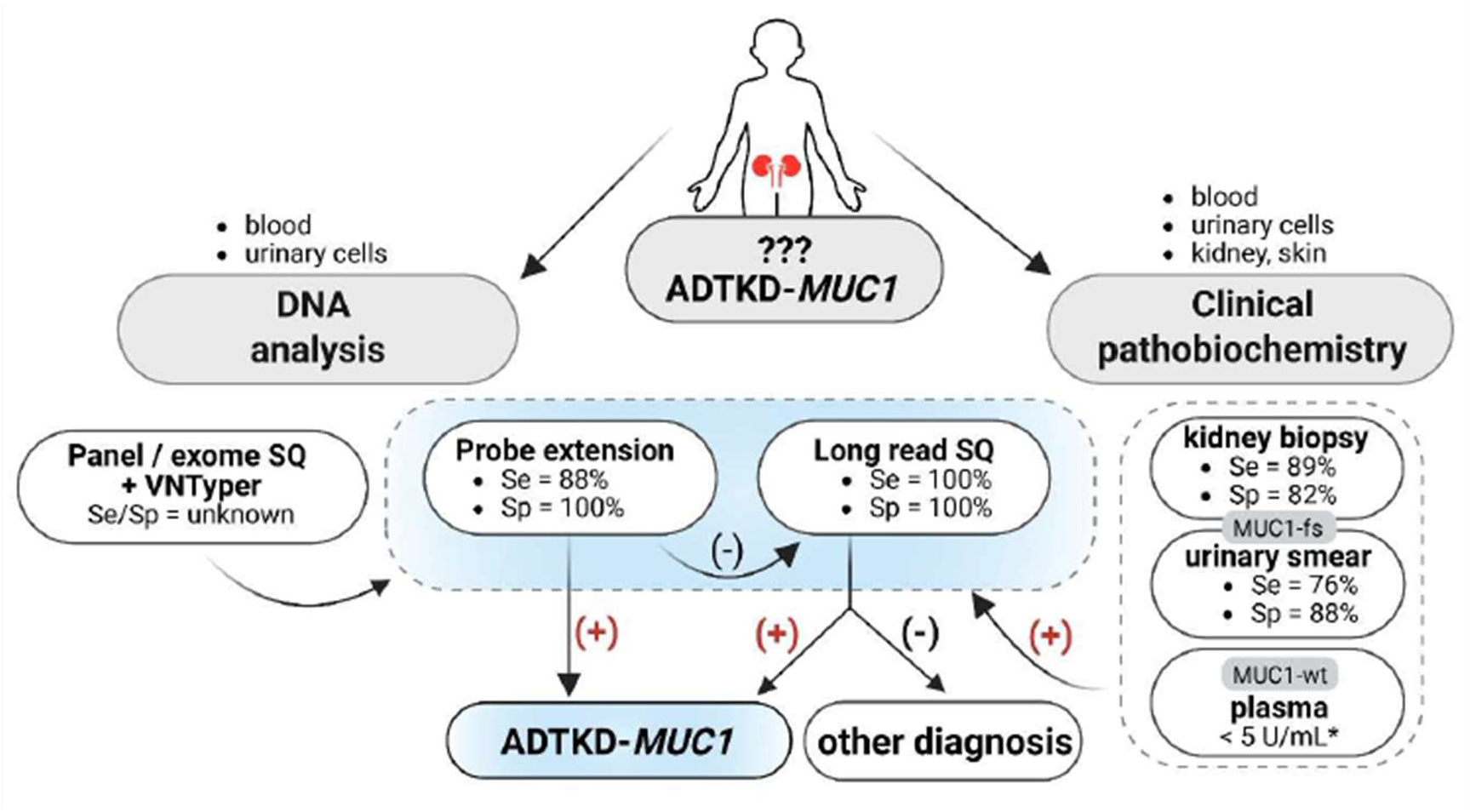
Diagnostic scheme for ADTKD-*MUC1*. The diagnosis of ADTKD-*MUC1* involves a combination of clinical evaluation, family history assessment, and specialized genetic, immunohistochemical and biochemical testing due to the unique challenges posed by the *MUC1* VNTR region. There are currently several methods available for *MUC1* genetic testing. In the United States, CLIA-approved genetic testing for ADTKD-ML/C1 primarily utilizes a mass spectrometry-based probe-extension assay, which has a sensitivity (Se) of approximatively 88%, as it detects only of a subset of known frameshift mutations, and a specificity (Sp) approaching 100%. This assay thus represents currently the first-choice method for diagnosing ADTKD-ML/C1. In cases with negative results, Single Molecule, Real-Time (SMRT) sequencing of *MUC1* may be performed at specialized research or diagnostic centers. This method offers sensitivity (Se) and specificity (Sp) approaching 100%. Compared to the probe-extension assay, SMRT sequencing enables exact mapping of the mutation within the full repeat sequence and allows for detailed sequence comparison with potential application in risk stratification and personalized clinical management of affected individuals. *MUC1* frameshift mutations have also been identified using short-read sequencing-based approaches in combination with specialized bioinformatics tools such as VNTyper. However, the sensitivity and specificity of these methods have not yet been fully evaluated, and their performance remains uncertain compared to probe-extension or long-read assays. Genetic diagnosis of ADTKD-ML/C1 may also be supported by immunodetection of MUClfs in kidney tissue or urinary cells as well as by the observation of markedly reduced plasma concentration of *MUC1*. Created with BioRender.com

In conclusion, we describe a robust SMRT sequencing-based methodology that enables detailed analysis of the *MUC1* VNTR sequence facilitating highly accurate diagnosis of ADTKD-*MUC1*. Our study reports extensive allelic and mutational variability within the *MUC1* VNTR, including numerous novel repeat sequences. These variants can be incorporated into a comprehensive *MUC1*-specific motif dictionary, which may improve the bioinformatics detection of *MUC1* mutation using short-read sequencing-based approaches [20, 30]. Importantly, we also demonstrate the potential prognostic value of individual- and family-specific VNTR structures, highlighting their possible use in risk stratification and personalized clinical management of affected individuals. We welcome collaboration and can provide both genetic testing and immunohistochemical screening as described in this study. For further information or to initiate testing, please see www.rikd.org/resources.

## Acknowledgements

We sincerely thank the many participating individuals and their families, together with their primary physicians and colleagues, whose generous contributions of clinical data and biological specimens made this work possible. Their commitment has been invaluable and continues to advance our understanding of ADTKD-*MUC1*. We would like to thank Anna Greka (Broad Institute of MIT and Harvard) for her comments on the manuscript.

## Author Contribution Statement

K. Kidd, M. živná, AJ. Bleyer and S.Kmoch conceived the concept, designed and coordinated the study. A. Vrbacká, A.Přistoupilová, V. Janoušek, M. Radina, P. Vyleťal, I. Bitar, V. Stránecký, L. Steiner-Mrázová, H. Trešlová, V. Barešová, D. Mušálková, J. Sovová, K. Hodaňová, H. Hartmannová, K. Svojšová, T. Kmochová, B. Blumenstiel, D. Toledo, M. DiStefano and M. DeFelice were responsible for acquisition and analysis of data. K. Kidd, A Taylor, L. Martin, A. Sanchez, and AJ. Bleyer organized, curated and maintained the Wake Forest ADTKD registry. R. Ryšavá, I. Lajtmannová, S. Bloudíčková-Rajnochová, O. Viklický, G. Papagregorius, C. Deltas, Ch. Stavrou, S. Jorge, JA Lopes, M. Rodrigues, E. Elhassan, M. Clince, C. Rowan, P. Conlon, O. Teltsh, GL. Cavalleri and AJ. Bleyer collected and curated clinical materials and associated patient information for research and registry purposes. AJ. Bleyer and S. Kmoch drafted the manuscript. All authors contributed to revision of the final version of the manuscript and approved the final submitted version.

## Data sharing statement

This is a clinical study, and individual-level clinical and genetic data cannot be shared to protect participant privacy. However, aggregated data on *MUC1* variation are provided in the Supplementary Material, and a periodically updated list of all identified VNTR repeats is available at: https://github.com/pristanna/muc1repeats.

