## Supplement for "Long-Read Sequencing of the *MUC1* VNTR: Genomic Variation, Mutational Landscape, and Its Impact on ADTKD Diagnosis and Progression"

The authors to give readers additional information about their work have provided this appendix.

**Table of contents**

**Supplementary methods** (p. 2-4)

**Supplementary references** (p. 4)

**Mass Spectrometry-Based Assay for the Molecular Diagnosis of ADTKD-MUC1**

A probe (primer) extension assay that in principle detects only mutations that are destroying MwoI restriction site within mutated VNTR was performed using a CLIA-approved mass spectrometry-based assay at the Broad Institute of MIT and Harvard (Cambridge, MA) [1; 2] . In this assay, the Mwol restriction endonuclease is used to cleave the 5′-GCNNNNN/NNGC-3′ recognition sequence within each of the 60-bp repeat units (where N represents any nucleotide). However, If a cytosine duplication expands a 7C homopolymer to 8Cs, only the mutated repeat escapes cleavage by the Mwol endonuclease and can be specifically amplified by PCR. The presense of the duplication in the PCR product is then interrogated through a DNA polymerase-mediated extension of a specific oligonucleotide probe. The molecular weight of the extended analyte is subsequently detected using matrix-assisted laser desorption/ionization time-of-flight (MALDI-TOF) mass spectrometry. Detection of the extended analyte at 5904 Da indicates the presence of an extra cytosine. In principle, the assay can also detect other types of frameshift mutations, such as the duplication of an adenosine adjacent to a 7C homopolymer, or the 58_59insG, which are characterized by the presence of an extended analytes at 6571 Da and 5944.85 Da, respectively [1].

**Long range PCR amplification and Long-read sequencing of MUC1**

Whole blood was collected and DNA isolated by standard methodology. The VNTR region of the MUC1 was PCR amplified as previously described 12. Each sample was amplified in 8 separate reaction tubes (25ul each), with the resulting PCR products (~1500 – 6000 bp long) pooled and purified using magnetic beads SPRI (Beckman Coulter, Inc., Brea, CA, USA) according to the manufacturer's protocol. Beads-purified PCR products (amplicons) were quantified using Qubit 2.0 Fluorometric Quantitation (Thermo Fisher Scientific, Waltham, MA, USA ) with QubitTM 1x dsDNA HS Assay Kit (Thermo Fisher Scientific, Waltham, MA, USA). Lengths of amplified MUC1 alleles were estimated using the 5200 Fragment Analyzer System (Agilent Technologies, Santa Clara, CA United States) on the HS Large Fragment Kit (DNF-493-0500) with 55 cm capillaries (A2300-1250-5580). Samples were sequenced on the Pacific Biosciences (Menlo Park, CA, USA) Sequel I system according to the manufacturer's protocol using the SMRTbell Express Template Prep Kit 2.0 and the Sequel Sequencing Kit 3.0 using SMRT Cell 1M v3 LR Tray.

**Bioinformatic analysis of sequence data**

To obtain highly accurate reads, Circular Consensus Sequence (CCS) analysis was performed using SMRT Link (version 8.0). The resulting CCS FASTQ files were analyzed using the proprietary bioinformatics pipeline, called PacMUC1. First, a reference sequence with 120 contigs was created, with each contig containing prerepeats (1, 2, 3, 4, 5) and afterrepeats (6, 7, 8, 9), separated by 1-120 canonical X repeats. The CCS FASTQ files were mapped to the reference sequence using bwa-mem (version 0.7.16a-r1181) and postprocessed using samtools (version 1.9) 29. Chimeric, incomplete, and low quality reads were removed, and the aligned sequences were displayed in the Integrative Genomics Viewer (IGV, version 2.16.2) for visual inspection. To determine the length of each of the two alleles for each sample, idxstats were generated using samtools (version 1.9) and peaks with the highest number of mapped reads were selected. To find changes from the reference sequence we used the deep neural network-based variant caller Clair (version 2.0.7), https://github.com/HKU-BAL/Clair and Clair3 (version 1.0.10), https://github.com/HKU-BAL/Clair3 with a model trained for PacBio data. The resulting VCF file and bcftools (version 1.10.2) were used to create a consensus sequence for each of the determined alleles. Custom Python scripts were used to separate the consensus sequences into individual repeats, classify them (each unique 60-base repeat segment is represented by a different letter or number, corresponding to the repeats previously identified by Kirby et al. [3] and Wenzel et al. [4], calculate statistics, and generate reports. The obtained MUC1 VNTR sequences are reported as a sequence of unique repetitive units, characterized by numbers (pre- and after- repeats) and letters (canonical repeats), as originally described by Kirby et al. 4. A minimum read coverage of 10× was required for reliable determination of the VNTR allele structure. In addition, for related individuals, only a single representative allele was reported to prevent artificially inflated allele occurrence or frequency. A minimum read coverage of 10× was required also for reliable identification of the frameshift mutation. Cases in which the mutation was clearly identified (eg. visible in the IGV) but supported by fewer than 10x reads were reported as inconclusive.

**Statistical Analysis**

Spearman’s rank correlation was used to assess the association between eGFR decline (calculated as eGFR – 120 / age) and the difference in repeat unit length between the wild-type allele and the frameshift mutation–harboring allele.

To further evaluate independent contributions of allele lengths, we performed linear regression with eGFR decline as the dependent variable. Predictor variables included wild-type (WT) allele length and mutated allele length. Standardized regression coefficients (β), standard errors (SE), p-values, and model $R^2$ were reported. Sequential model testing was used to assess incremental contributions of predictors. All statistical analyses were conducted in R, with two-tailed significance set at $p < 0.05$.

1 2 3 4 5 X D E C F X X A B X E C X X X A A B X X X X X X G A B X X X X X X X V 6' 7 8 9
1 2 3 4 5 aJ X D E C X X X A A B D E C X aF aI A A B D E X A A B X X X X A A B aD E C X X X A A B X X X X X X A A B X X X X X X X V 6' 7 8 9
1 2 3 4 5 C aN D E C F X X A B D E C X X X A A B X X X X X X X X X G A B X X X X X X X V 6' 7 8 9
1 2 3 4 5 C X aK E C F X X A B D E C X X X A A B X X X X X X G A B X X X X X X X V 6' 7 8 9
1 2 3 4 5 C X aK E C F X X A B D E C X X X A A B X X X X X X G A B X X X X X X X V 6+ 7 8 9
1 2 3 4 5 C X D C T X A B D E C X X X A G B X X X X X X X X X E C X X X A A B X X X X X X X X X I X A J B X X X X X X X V 6' 7 8 9
1 2 3 4' 5 C X D C X X X A A B X X X X B A A B B X X X X X X G A B X X X X X X X X V 6 7 8 9
1 2 3 4' 5 C X D C X X X A A B X X X X B A A B X X X X X X X X G A B B X X X X X V 6 7 8 9
1 2 3 4' 5 C X D C X X X A A B X X X X B A A B X X X X X X X X G A B X X X X X X X X 6 aO 8 9
1 2 3 4' 5 C X D C X X X A A B X X X X B A A B X X X X X X X X G A B X X X X X X X X V 6 7 8 9
1 2 3 4' 5 C X D C X X X A A B X X X X B A A B X X X X X X X X G A B X X X X X X X X X 6 7 8 9
1 2 3 4' 5 C X D C X X X A A C C X D C X X X A A B X X X X B A A B X X X X X X X X X V 6 7 8 9
1 2 3 4' 5 C X D C X X X:59dupC A A B X X X X B A A B X X X X X X X X G A B B X X X X X V 6 7 8 9
1 2 3 4 5 C X D E aX F X X X A A B X X X X X X X G A B X X X X X X X V 6' 7 8 9
1 2 3 4 5 C X D E C F bC X A B D E C X X X A A B X X X X X X G A B X X X X X X G A B B X X X X X X V 6' 7 8 9
1 2 3 4 5 C X D E C F X X A A B D E C X X X A A B X X X X X X X G A B X X X X V 6+ 7 8 9
1 2 3 4 5 C X D E C F X X A B D E C Q X X A A B X X X X X X X G A B X X X X X X X V 6' 7 8 9
1 2 3 4 5 C X D E C F X X A B D E C X B X A A B X X X X X X X G A N R X X X G A B X X X X X X X V 6' 7 8 9
1 2 3 4 5 C X D E C F X X A B D E C X X W A A B X X G A B X X X X X X X V 6' 7 8 9
1 2 3 4 5 C X D E C F X X A B D E C X X W A A B X X G A B X X X X X X X X X X V 6' 7 8 9
1 2 3 4 5 C X D E C F X X A B D E C X X W A A B X X X X X X G A B X X X X X X X V 6' 7 8 9
1 2 3 4 5 C X D E C F X X A B D E C X X W A A B X X X X X X X G A B X X X X X X X V 6' aS 8 9
1 2 3 4 5 C X D E C F X X A B D E C X X W A G B X X X X X G A B X X X X X X X X V 6' 7 8 9
1 2 3 4 5 C X D E C F X X A B D E C X X W A G B X X X X X G bG B X X X X X X X X V 6' 7 8 9
1 2 3 4 5 C X D E C F X X A B D E C X X W A G B X X X X X X G A B X X X X X X X V 6' 7 8 9
1 2 3 4 5 C X D E C F X X A B D E C X X X A A B B X X X X X X G A B X X X X X X X V 6' 7 8 9
1 2 3 4 5 C X D E C F X X A B D E C X X X A A B X B X X X X X G A B X X X X X X X X X V 6' 7 8 9
1 2 3 4 5 C X D E C F X X A B D E C X X X A A B X X X X X G A B X X X X X X X V 6' 7 8 9
1 2 3 4 5 C X D E C F X X A B D E C X X X A A B X X X X X X G A B X X X X X X G A B B X X X X X X V 6' 7 8 9
1 2 3 4 5 C X D E C F X X A B D E C X X X A A B X X X X X X G A B X X X X X X X V 6' 7 8 9
1 2 3 4 5 C X D E C F X X A B D E C X X X A A B X X X X X X X G A B X X X aR G A B X X X X X X X V 6' 7 8 9
1 2 3 4 5 C X D E C F X X A B D E C X X X A A B X X X X X X X G A B X X X X X G A B X X X X X G B V 6' 7 8 9
1 2 3 4 5 C X D E C F X X A B D E C X X X A A B X X X X X X X G A B X X X X X G A B X X X X X X X V 6' 7 8 9
1 2 3 4 5 C X D E C F X X A B D E C X X X A A B X X X X X X X G A B X X X X X G A B X X X X X X X V 6+ 7 8
1 2 3 4 5 C X D E C F X X A B D E C X X X A A B X X X X X X X G A B X X X X X G A B X X X X X X X V 6+ 7 8 9
1 2 3 4 5 C X D E C F X X A B D E C X X X A A B X X X X X X X G A B X X X X X X V 6' 7 8 9
1 2 3 4 5 C X D E C F X X A B D E C X X X A A B X X X X X X X G A B X X X X X X X V 6' 7 8 9
1 2 3 4 5 C X D E C F X X A B D E C X X X A A B X X X X X X X G A B X X X X X X X V 6+ 7 8 9
1 2 3 4 5 C X D E C F X X A B D E C X X X A A B X X X X X X X G A B X X X X X X X V 6+ 7 8 9
1 2 3 4 5 C X D E C F X X A B D E C X X X A A B X X X X X X X G A N R X X X G A B X X X X X X V 6' 7 8 9
1 2 3 4 5 C X D E C F X X A B D E C X X X A A B X X X X X X X G A N R X X X G A B X X X X X X X V 6' 7 8 9
1 2 3 4 5 C X D E C F X X A B D E C X X X A A B X X X X X X X G G B X X X X G A B X X X X X X X V 6' 7 8 9
1 2 3 4 5 C X D E C F X X A B D E C X X X A A B X X X X X X X X G A B X X X X X X X V 6' 7 8 9
1 2 3 4 5 C X D E C F X X A B D E C X X X A A B X X X X X X X X V 6' 7 8 9
1 2 3 4 5 C X D E C F X X A B D E C X X X A B X X X X X X G A B X X X X X X X V 6' 7 8 9
1 2 3 4 5 C X D E C F X X A B D E C X X X A B X X X X X X X G A B X X X X X X X V 6' 7 8 9
1 2 3 4 5 C X D E C F X X A B D E C X X X G A B X X X X X X V 6' 7 8 9

**Pre-repeats units**

AAGGAGACTTCGGCTACCCAGAGAAGTTCAGTGCCCAGCTCTACTGAGAAGAATGCTGTG 1
AGTATGACCAGCAGCGTACTCTCCAGCCACAGCCCCGGTTCAGGCTCCTCCACCACTCAG 2
GGACAGGATGTCACTCTGGCCCCGGCCACGGAACCAGCTTCAGGTTCAGCTGCCACCTGG 3
GGACAGGATGTCACCTCGGTCCCAGTCACCAGGCCAGCCCTGGGCTCCACCACCCCGCCA 4
GGACAGGATGTCACCTCGGTCCCAGTCACCAGGCCAGCCCTGGGCTCCACCACCCCACCA 4'
GGACAGGATGTCACCTCCGTCCCAGTCACCAGGCCAGCCCTGGGCTCCACCACCCCGCCA 4''
GGACAGGATGTCACCTCGGTCCCAGTCACCAGGCCAGCACTGGGCTCCACCACCCCGCCA 4''
GCCCACGATGTCACCTCAGCCCCGGACAACAAGCCAGCCCCGGGCTCCACCGCCCCCCCA 5
GCCCACGATGTCACCTCAGCCCCGGACACCAGGCCGGCCCCGGGCTCCACCGCCCCCCAA 5C

KETSATQRSSVPSSTEKNAV 1
SMTSSVLSSHSPGSGSSTTQ 2
GQDVTLAPATEPASGSAATW 3
GQDVTSVPVTRPALGSTTPP 4
GQDVTSVPVTRPALGSTTPP 4'
GQDVTSVPVTRPALGSTTPP 4''
GQDVTSVPVTRPALGSTTPP 4'''
AHDVTSAPDNKPAPGSTAPP 5
AHDVTSAPDTRPAPGSTAPQ 5C

**After-repeats units**

GCCCACGGTGTCACCTCGGCCCCGGACACCAGGCGGGCCCCGGGCTCCACCCCGGCCCCG 6
GCCCACGGTGTCACCTCGGCCCCGGACACCAGGCCGGCCCCGGGCTCCACCCCGGCCCCG 6'
GGCTCCACCGCCCCCCCAGCCCACGGTGTCACCTCGGCCCCGGACACCAGGCCGGCCCCG 7
GGCTCCACCGCCTCCCCAGCCCACGGTGTCACCTCGGCCCCGGACACCAGGCCGGCCCCG 7'
GGCTCCACCGCCCCCCCAGCCCATGGTGTCACCTCGGCCCCGGACAACAGGCCCGCCTTG 8
GGCTCCACCGCCCCTCCAGTCCACAATGTCACCTCGGCCTCAGGCTCTGCATCAGGCTCA 9

AHGVTSAPDTRRAPGSTPAP 6
AHGVTSAPDTRPAPGSTPAP 6'
GSTAPPAHGVTSAPDTRPAP 7
GSTASPAHGVTSAPDTRPAP 7'
GSTAPPAHGVTSAPDNRPAL 8
GSTAPPVHNVTSASGSASGS 9

Types  of frameshift mutations identified
18_31delGGCCCCGGACACCA
23dupC
42_57dupGGGCTCCACCGCCCCC
54_55insA
54_55insG
56_59dupCCCC
58_59insG
59dupC
60dupA

wt
GCCCACGGTGTCACCTCGGCCCCGGACACCAGGCCGGCCC*CGGGCTCC* ***A*** *CCG* *CCCCCCC* AGCCCACGGTGTCACCTCGGCCCCGGACACCAGGCCGGCCCCGGGCTCCACCGCCCCCCCAGCCCACGGTGTCACCTCGGCCCCGGACACCAGGCCGGCCCCGGGCTCCACCGCCCCCCCA

pre-VNTR
c.326_350dup, p.(Ser119Profs∗119 de Haan, KI 2023

types o

Mutated VNTR alleles

1 2 3 4' 5 C X D C X X X:59dupC A A B X X X X B A A B X X X X X X X G A B B X X X X V 6 7 8 9
1 2 3 4 5 C X D E C F X X A B D E:23dupC C X X X A A B X X X X X X G A B X X X X X X V 6' 7 8 9
1 2 3 4 5 C X D E C F X X A B:59dupC D E C X X X A A B X X X X X G A B X X X X X X V 6' 7 8 9
1 2 3 4 5 C X D E C F X X A B:59dupC X E C X X X A A B X X X X X X G A B X X X X X X V 6' 7 8 9
1 2 3 4 5 C X D E C F X:59dupC X A B D E C X X X A A B X X X X X X G A N R X X X G A B X X X X X X V 6' 7 8 9
1 2 3 4 5 C X D E C X H X A A B D E C X aH:54_55insA X A A B X X X X X X X X X X X E C X X X G A B X X X X X X X X X X V V I G J B B X X X X V 6' 7' 8 9
1 2 3 4 5 C X D E C X H X A A B:59dupC D E C X X X A A B X X X X X E C X X X A A B X X X X X X X X X X X X X X X V X A J B X X X X X X V 6' 7 8 9
1 2 3 4 5 C X D E C X T X A B D E C X X X A A B X B X X X X X G bJ:54_55insG X X X X X A A B X X X X X I X A J B X X X X X X X V 6' 7 8 9
1 2 3 4 5 C X D E C X T X A B D E C X X X A G B X X X X X X X X X E C:60dupA X X X A A B X X X X X X X X X X I X A J B X X X I X A J B X X X X X X V 6' 7 8 9
1 2 3 4 5 C X D E C X X A A B X X X:59dupC X X A A A B X X X X X A A X X X X X X X X A A B X X X X B B X X A A B X X X X B A A X X X X X X X X A A B X X X X X X V 6 7 8 9
1 2 3 4 5 C X D E C X X X A A B B X X X X A A A B X X X X:59dupC X A A X X X X X X X X X A A B X X X X B B X X A A B X X X X B A A X X X X X G A B X X X X V 6 7 8 9
1 2 3 4 5 C X D E C X X X A A B B X X X X A A A X X X X X X A A X X X X X X X X A:23dupC A B X X X X B X X A A B X X X X B A A X X X X X G A B B X X X X X X V 6 7 8 9
1 2 3 4 5 C X D E C X X X A A B X X X X A A A B X X X X X A A X X X X X X X X A A B X X X X:59dupC B B X X A A B X X X X B A A X X X X X X X G A B X X X X X X X V 6 7 8 9
1 2 3 4 5 C X D E C X X X A A B X X X X A A A B X X X X X A A X X X X X X X X A A B X X X:59dupC X B B X X A A B X X X X B A A X X X X X X X G A B X X X X X X X V 6 7 8 9
1 2 3 4 5 C X D E C X X X A A B X X X X X A A A B X X X X A A X X X X X X X X A A B X X X:59dupC X B B X X A A B X X X X B A A X X X X X G A B X X X X V 6 7 8 9
1 2 3 4 5 C X D E C X X X A A B X X X X X A A A B X X X X X A A X X X X X X X B B X X:58_59insG A B X X X X X A A X X X X X X X B B X X A A B X X X X B A A X X X X X X X G A B X X X X X X X V 6 7 8 9
1 2 3 4 5 C X D E C X X X A A B X X X X X A A A B X X X X X A A X X X X X X X G B:58_59insG A B X X X X B X X A A B X X X X B A A X X X X X G A B X X X X X X X V 6 7 8 9
1 2 3 4 5 C X D E C X X X A A B X X X X X A A A B X X X X X A A X X X X X X X X A A B X X X X B B X:60dupA X A A B X X X X B A A X X X X X X X G A B X X X X X X X V 6 7 8 9
1 2 3 4 5 C X D E C X X X A A B X X X X X A A A B X X X X X A A X X X X X X X X A A B X X X X B:59dupC X X A A B X X X X B A A X X X X X G A B X X X X X X X V 6 7 8 9
1 2 3 4 5 C X D E C X X X A A B X X X X X A A A B X X X X X A A X X X X X X X X A A B X X:59dupC X X B X X A A B X X X X B A A X X X X X G A B X X X X X X X V 6 7 8 9
1 2 3 4 5 C X D E C X X X A A B X X X X X A A A B X X X X X A A X X X X X X:59dupC X A A B X X X X B X X A A B X X X X X A A X X X X B G A B X X X X X X X V 6 7 8 9
1 2 3 4 5 C X D E C X X X A A B X X X X X A A A B X X X X X A A X:59dupC X X X X X X A A B X X X X B X X A A B X X X X B A A X X X X X G A B X X X X X X X V 6 7 8 9
1 2 3 4 5 C X D E C X X X A A B X X X X X A A A B X X X X X X X X X X:59dupC X A A B X X X X B X X A A B X X X X B A A X X X X X G A B X X X X X X X V 6 7 8 9
1 2 3 4 5 C X D E C X X X A A B X X X X X A A A B X X X:59dupC X X A A X X X X X X X X X A A B X X X B X X A A B X X X X B A A X X X X X G A B X X X X X X X V 6 7 8 9
1 2 3 4 5 C X D E C X X X A A B X X X X X A A A B X X X:59dupC X X A A X X X X X X X X X A A B X X X B X X A A B X X X X B A A X X X X X G A B X X X X X X X V 6 7 8 9
1 2 3 4'' 5 C X D E C X X X A A B X X X X X A A A B X X X:59dupC X X A A X X X X X X X X X A A B X X X X B X X A A B X X X X B A A X X aF X X G A B X X X X X X X V 6 7 8 9
1 2 3 4 5 C X D E C X X X A A B X X X X X A A A B X:59dupC X X X X A A X X X X X X X X X A A B X X X X B X X A A B X X X X X A A X X X X G A B X X X X X X X V 6 7 8 9
1 2 3 4 5 C X D E C X X X A A B X X X X X A A A B:59dupC X X X X X A A X X X X X X X X X A A B X X X X B B X X A A B X X X B A A X X X X X X X G A B X X X X X X X V 6 7 8 9
1 2 3 4 5 C X D E C X X X A A B X X X X:59dupC A A A B X X F X X A A X X X X X X X A A B X X X X B X X A A B X X X B A A X X X X X G A B X X X X X X V 6 7 8 9
1 2 3 4 5 C X D E C X X X A A B X X X X:59dupC A A A B X X X X X A A X X X X X X X X A A B X X X X B B X X A A B X X X X B A A X X X X X X X G A B X X X X X X X V 6 7 8 9
1 2 3 4 5 C X D E C X X X A A B X X X X:59dupC X A A A B X X X X X A A X X X X X X X X A A B X X X X B X X A A B X X X X B A A X X X X B G A B X X X X X X X V 6 7 8 9
1 2 3 4 5 C X D E C X X X A A B X X X X:59dupC X A A A B X X X X X A A X X X X X X X X A A B X X X X B X X A A B X X X X B A A X X X X B G A B X X X X X X X X 6 7 8 9
1 2 3 4 5 C X D E C X X X A A B X X:59dupC X X X A A A B X X X X X X X X X X X X X A A X X X X X B X X A A B X X X X B A A X X X X G A B X X X X X X X X V 6 7 8 9
1 2 3 4'' 5 C X D E C X X X A A B X:59dupC X X X X A A A B X X X X X A A X X X X X X X X A A B X X X X B X X A A B X X X B A A X X X X X G A B X X X X X X X V 6 7 8 9
1 2 3 4 5 C X D E C X X X A A B:59dupC X X X X X A A A B X X X X X A A X X X X X X X A A B X X X X B X X A A B X X X B A A X X X X X G A B X X X X X X V 6 7 8 9
1 2 3 4 5 C X D E C X X X:59dupC A A B X X X X X A A A B X X X X X A A X X X X X X X A A B X X X X B B X X A A B X X X X B A A X X X X X X X G A B X X X X X X X V 6 7 8 9
1 2 3 4 5 C X D E C X X:56_59dupCCCC X A B X X X X X A A A B X X X X X X X X X X X X A A B X X X X B X X A A B X X X X B A A X X X X X G A B X X X X X X X V 6 7 8 9
1 2 3 4 5 C X D E C X X:59dupC X A A B X X X X X A A A B X X X X X X X X X X X X A A B X X X X B X X A A B X X X X B A A X X X X X G A B X X X X X X X V 6 7 8 9
1 2 3 4 5 C X D E C X X:59dupC X A A B X X X X X A A B X X X X X A A X X X X X X X X A A B X X X X B B X A A B X X X X B A A X X X X X X X X A A B X X X X X X X V 6 7 8 9
1 2 3 4 5 C X D E C X:59dupC bI X A A B X X X X X A A A B X X X X X A A X X X X X X X X A A B X X X X B B X X A A B X X X X B A aT X X X X X X X G A B X X X X X X X V 6 7 8 9
1 2 3 4'' 5 C X D E C X:59dupC X X A A B X X X X X A A A B X X X X X A A X X X X X X X X A A B X X X X B X X A A B X X X X B A A X X X X X G A B X X X X X X X V 6 7 8 9

1 2 3 4 5 C X D E C:42_57dupGGGCTCCACCGCCCCC X X X A A B X X X X X A A A B X X X X X X B B X X A A B X X X X X A A B X X X X X G A B X X X X X X X V 6 7 8 9
1 2 3 4' 5 C X D:59dupC C X X X A A B X X X X B A A B X X X X bE X G A B X X X X X V 6 7 8 9
1 2 3 4 5 C X D:59dupC E C F X X A B X E C X X X A A B X X X X X X G A B X X X X X X V 6' 7 8 9
1 2 3 4 5 C X D:59dupC E C X T X A B D E C X X X A A B X X X X X X X X X X I X A J B X X X X X X V 6' 7 8 9
1 2 3 4 5 C X:59dupC D E C F X X A B D E C X X X A A B X X X X X X G A B X X X X X X V 6' 7 8 9
1 2 3 4' 5:59dupC C X D C X X X A A B X X X X B A A B X X X X X X X G A B X X X X X X X V 6 7 8 9
1 2 3 4 5:59dupC C X D E C F X X A B D E C X X X A A B X X X X X G A B X X X X X X V 6' 7 8 9
1 2 3 4 5:59dupC C X D E C F X X A B D E C X X X A A B X X X X X X G A B X X X X X X V 6' 7 8 9
1 2 3 4 5:59dupC C X D E C F X X A B X E C X X X A A B X X X X X X G A B X X X X X X V 6' 7 8 9
1 2 3 4' 5:59dupC C X D X A A A A B X X X X B A A B X X X X X X X G A B X X X X X X X X 6 7 8 9
1 2 3 4 5 C X D E C F X X:18_31delGGCCCCGGACACCA A B X E C X X X A A B X X X X X X G A B X X X X X X V 6' 7 8 9

| Gender | eGFR-120/ | Fast = 1/Slo | WT allele le | Mutated al | # of wild ty | # of mutated repeats | wt-mt |
|---|---|---|---|---|---|---|---|
| Female | -6,47059 | 1 | 34 | 68 | 31 | 37 | -34 |
| Male | -6,11111 | 1 | 34 | 70 | 23 | 47 | -36 |
| Female | -5 | 1 | 71 | 68 | 31 | 37 | 3 |
| Male | -4,78261 | 1 | 61 | 35 | 3 | 32 | 26 |
| male | -4,58333 | 1 | 35 | 74 | 24 | 50 | -39 |
| Female | -4,58333 | 1 | 71 | 71 | 28 | 43 | 0 |
| Female | -4,58333 | 1 | 35 | 70 | 23 | 47 | -35 |
| Female | -4,58333 | 1 | 66 | 70 | 23 | 47 | -4 |
| Female | -4,23077 | 1 | 34 | 71 | 23 | 49 | -37 |
| Female | -4,07407 | 1 | 72 | 67 | 10 | 57 | 5 |
| Female | -4,07407 | 1 | 84 | 69 | 24 | 46 | 15 |
| Female | -4,07407 | 1 | 35 | 71 | 36 | 35 | -36 |
| male | -4,07407 | 1 | 33 | 57 | 5 | 52 | -24 |
| Male | -3,92857 | 1 | 35 | 66 | 7 | 59 | -31 |
| female | -3,92857 | 1 | 35 | 70 | 20 | 50 | -35 |
| Female | -3,92857 | 1 | 36 | 59 | 11 | 48 | -23 |
| Female | -3,92857 | 1 | 36 | 71 | 13 | 58 | -35 |
| Female | -3,92857 | 1 | 71 | 70 | 12 | 58 | 1 |
| Female | -3,92857 | 1 | 35 | 71 | 36 | 35 | -36 |
| Female | -3,92857 | 1 | 35 | 69 | 30 | 39 | -34 |
| Male | -3,7931 | 1 | 37 | 71 | 29 | 42 | -34 |
| Female | -3,66667 | 1 | 37 | 71 | 29 | 42 | -34 |
| Male | -3,66667 | 1 | 30 | 67 | 16 | 51 | -37 |
| Male | -3,66667 | 1 | 41 | 74 | 16 | 58 | -33 |
| Male | -3,63467 | 1 | 69 | 69 | | | 0 |
| male | -3,54839 | 1 | 25 | 42 | 7 | 35 | -17 |
| Male | -3,54839 | 1 | 35 | 35 | 0 | 36 | 0 |
| Male | -3,54839 | 1 | 71 | 35 | 0 | 35 | 36 |
| Female | -3,54839 | 1 | 68 | 35 | 0 | 35 | 33 |
| Female | -3,33983 | 1 | 35 | 67 | 7 | 60 | -32 |
| Male | -3,33333 | 1 | 34 | 68 | 31 | 37 | -34 |
| Male | -3,33333 | 1 | 77 | 72 | 20 | 52 | 5 |
| female | -3,23529 | 1 | 36 | 69 | 13 | 56 | -33 |
| Male | -3,23529 | 1 | 35 | 70 | 23 | 47 | -35 |
| Male | -3,23529 | 1 | 35 | 71 | 15 | 56 | -36 |
| Female | -3,19048 | 1 | 35 | 35 | 0 | 35 | 0 |
| male | -3,15455 | 1 | 74 | 71 | 34 | 37 | 3 |
| female | -3,14286 | 1 | 34 | 57 | 5 | 52 | -23 |
| Male | -3,14286 | 1 | 28 | 34 | 0 | 34 | -6 |
| Female | -3,12444 | 1 | 72 | 42 | 7 | 35 | 30 |
| Female | -3,05556 | 1 | 35 | 64 | 29 | 35 | -29 |
| female | -3,05143 | 1 | 69 | 69 | 13 | 56 | 0 |
| Female | -2,97297 | 1 | 74 | 66 | 7 | 59 | 8 |
| female | -2,96944 | 1 | 74 | 74 | 24 | 50 | 0 |
| Male | -2,89847 | 1 | 42 | 74 | 16 | 58 | -32 |
| Female | -2,89474 | 1 | 35 | 35 | 0 | 35 | 0 |
| Male | -2,89474 | 1 | 25 | 70 | 12 | 58 | -45 |
| Female | -2,82051 | 1 | 69 | 69 | 31 | 38 | 0 |
| female | -2,82051 | 1 | 35 | 34 | 9 | 25 | 1 |
| Female | -2,82051 | 1 | 71 | 41 | 3 | 38 | 30 |
| Female | -2,82051 | 1 | 35 | 70 | 15 | 56 | -35 |
| Female | -2,82051 | 1 | 55 | 35 | 0 | 35 | 20 |
| Female | -2,82051 | 1 | 74 | 86 | 38 | 48 | -12 |
| female | -2,77 | 1 | 70 | 71 | 15 | 56 | -1 |
| Female | -2,75 | 1 | 41 | 74 | 45 | 29 | -33 |
| Female | -2,75 | 1 | 67 | 35 | 2 | 33 | 32 |
| Male | -2,75 | 1 | 35 | 35 | 12 | 23 | 0 |
| Female | -2,75 | 1 | 42 | 70 | 12 | 58 | -28 |
| female | -2,74368 | 1 | 34 | 71 | 15 | 56 | -37 |
| Male | -2,68293 | 1 | 42 | 59 | 11 | 48 | -17 |
| Male | -2,68293 | 1 | 58 | 71 | 29 | 42 | -13 |
| Female | -2,68293 | 1 | 65 | 67 | 13 | 54 | -2 |
| Male | -2,68293 | 1 | 35 | 54 | 28 | 26 | -19 |
| female | -2,655 | 1 | 66 | 71 | 15 | 56 | -5 |
| Female | -2,55814 | 1 | 74 | 35 | 8 | 27 | 39 |
| Female | -2,55814 | 1 | 66 | 36 | 0 | 36 | 30 |
| Female | -2,52499 | 1 | 33 | 71 | 36 | 35 | -38 |
| Female | -2,5 | 1 | 37 | 35 | 12 | 23 | 2 |
| Female | -2,44444 | 0 | 34 | 71 | 29 | 42 | -37 |
| Female | -2,44444 | 0 | 35 | 35 | 0 | 35 | 0 |
| male | -2,3913 | 0 | 71 | 71 | 15 | 56 | 0 |
| Female | -2,3913 | 0 | 24 | 59 | 11 | 48 | -35 |
| Male | -2,3913 | 0 | 35 | 35 | 1 | 34 | 0 |
| Female | -2,34043 | 0 | | 41 | 3 | 38 | |
| Male | -2,29602 | 0 | 61 | 35 | 3 | 32 | 26 |
| Female | -2,29167 | 0 | 24 | 34 | 0 | 34 | -10 |
| Female | -2,2449 | 0 | | 35 | 2 | 33 | |
| Male | -2,2 | 0 | 67 | 35 | 2 | 33 | 32 |
| Male | -2,18027 | 0 | | 35 | 0 | 35 | |
| Male | -2,15686 | 0 | 74 | 35 | 0 | 35 | 39 |
| Female | -2,15686 | 0 | 42 | 86 | 38 | 48 | -44 |
| Female | -2,10145 | 0 | 71 | 34 | 7 | 27 | 37 |
| Female | -2,03704 | 0 | 69 | 74 | 40 | 34 | -5 |
| Female | -2,03704 | 0 | 75 | 60 | 16 | 44 | 15 |
| Male | -2,02243 | 0 | 34 | 67 | 7 | 60 | -33 |
| Male | -1,96898 | 0 | 74 | 67 | 7 | 60 | 7 |
| Male | -1,96785 | 0 | 73 | 67 | 7 | 60 | 6 |
| Male | -1,96429 | 0 | 78 | 71 | 40 | 31 | 7 |

| | WT allele le | Mutated al | # of wild ty | # of mutated repeats | |
|---|---|---|---|---|---|
| median | | | | | |
| fast | 37 | 69 | 15 | 46 | -15 |
| slow | 57 | 59 | 9 | 35 | 0 |

| | | | | | | | |
|---|---|---|---|---|---|---|---|
| Female | -1,93463 | 0 | 69 | 73 | 8 | 65 | -4 |
| Male | -1,92982 | 0 | 31 | 34 | 10 | 24 | -3 |
| Male | -1,92982 | 0 | 55 | 68 | 40 | 28 | -13 |
| Male | -1,92982 | 0 | 74 | 35 | 0 | 35 | 39 |
| Male | -1,92982 | 0 | 55 | 33 | 3 | 30 | 22 |
| Male | -1,91488 | 0 | 34 | 67 | 7 | 60 | -33 |
| Female | -1,89655 | 0 | 57 | 59 | 11 | 48 | -2 |
| Male | -1,89655 | 0 | 42 | 70 | 12 | 58 | -28 |
| Female | -1,89655 | 0 | 71 | 68 | 40 | 28 | 3 |
| Male | -1,86813 | 0 | 35 | 35 | 3 | 32 | 0 |
| Female | -1,86098 | 0 | 74 | 34 | 0 | 34 | 40 |
| Male | -1,83388 | 0 | 69 | 70 | 23 | 47 | -1 |
| Female | -1,83333 | 0 | 35 | 68 | 32 | 36 | -33 |
| Male | -1,83333 | 0 | 24 | 59 | 11 | 48 | -35 |
| Female | -1,83333 | 0 | 52 | 72 | 40 | 32 | -20 |
| Female | -1,83333 | 0 | 74 | 34 | 7 | 27 | 40 |
| Female | -1,83333 | 0 | 36 | 60 | 16 | 44 | -24 |
| Female | -1,80328 | 0 | | 59 | 11 | 48 | |
| Female | -1,72128 | 0 | 72 | 42 | 7 | 35 | 30 |
| Female | -1,71875 | 0 | 68 | 72 | 7 | 65 | -4 |
| Female | -1,69632 | 0 | 71 | 71 | 40 | 31 | 0 |
| Male | -1,69231 | 0 | 57 | 59 | 11 | 48 | -2 |
| Female | -1,69231 | 0 | 62 | 35 | 10 | 25 | 27 |
| Female | -1,61963 | 0 | 78 | 71 | 40 | 31 | 7 |
| Male | -1,57143 | 0 | 71 | 35 | 1 | 34 | 36 |
| Female | -1,55554 | 0 | 35 | 35 | 2 | 33 | 0 |
| Male | -1,54325 | 0 | 76 | 35 | 2 | 33 | 41 |
| Male | -1,50211 | 0 | 74 | 86 | 38 | 48 | -12 |
| Male | -1,48649 | 0 | 41 | 68 | 40 | 28 | -27 |
| Female | -1,48649 | 0 | 41 | 68 | 40 | 28 | -27 |
| Male | -1,44977 | 0 | 35 | 35 | 12 | 23 | 0 |
| Female | -1,41932 | 0 | 35 | 72 | 7 | 65 | -37 |
| Female | -1,39653 | 0 | 68 | 34 | 7 | 27 | 34 |
| Male | -1,38489 | 0 | 27 | 71 | 6 | 65 | -44 |
| Female | -1,35893 | 0 | 74 | 59 | 11 | 48 | 15 |
| Female | -1,34189 | 0 | 35 | 35 | 2 | 33 | 0 |
| Male | -1,33703 | 0 | 71 | 73 | 41 | 32 | -2 |
| male | -1,33065 | 0 | 42 | 34 | 9 | 25 | 8 |
| Female | -1,26778 | 0 | 36 | 59 | 11 | 48 | -23 |
| Male | -1,25958 | 0 | 41 | 74 | 45 | 26 | -33 |
| Female | -1,2381 | 0 | 60 | 68 | 40 | 28 | -8 |
| Female | -1,22857 | 0 | 70 | 74 | 6 | 68 | -4 |
| Male | -1,21463 | 0 | 55 | 68 | 40 | 28 | -13 |
| Male | -1,19922 | 0 | 36 | 59 | 11 | 48 | -23 |
| female | -1,04515 | 0 | 42 | 34 | 9 | 25 | 8 |
| male | -1,01061 | 0 | 70 | 42 | 7 | 35 | 28 |
| Male | -0,92775 | 0 | 36 | 35 | 2 | 33 | 1 |
| female | -0,88163 | 0 | 35 | 74 | 6 | 60 | -39 |
| Male | -0,87286 | 0 | 36 | 59 | 11 | 48 | -23 |
| Male | -0,86909 | 0 | 60 | 73 | 41 | 32 | -13 |
| Male | -0,85027 | 0 | 73 | 72 | 7 | 65 | 1 |
| Female | -0,83957 | 0 | 71 | 35 | 0 | 35 | 36 |
| Female | -0,82458 | 0 | 74 | 34 | 0 | 34 | 40 |
| Female | -0,82272 | 0 | 73 | 35 | 10 | 25 | 38 |
| Male | -0,67543 | 0 | 73 | 71 | 43 | 28 | 2 |
| Male | -0,64568 | 0 | 35 | 35 | 2 | 33 | 0 |
| Male | -0,54791 | 0 | 68 | 60 | 16 | 44 | 8 |
| male | -0,40242 | 0 | 35 | 71 | 15 | 56 | -36 |
| female | -0,38907 | 0 | 70 | 42 | 7 | 35 | 28 |
| male | -0,31921 | 0 | 42 | 74 | 6 | 68 | -32 |
| female | 0,2504 | 0 | 33 | 57 | 5 | 52 | -24 |
